# Quantifying Induced Dipole Effects in Small Molecule Permeation in a Model Phospholipid Bilayer

**DOI:** 10.1101/2024.03.12.584668

**Authors:** Julia M. Montgomery, Justin A. Lemkul

## Abstract

The cell membrane functions as a semi-permeable barrier that governs the transport of materials into and out of cells. The bilayer features a distinct dielectric gradient due to the amphiphilic nature of its lipid components. This gradient influences various aspects of small molecule permeation and the folding and functioning of membrane proteins. Here, we employ polarizable molecular dynamics simulations to elucidate the impact of the electronic environment on the permeation process. We simulated eight distinct amino acid sidechain analogs within a 1-palmitoyl-2-oleoyl-*sn*-glycero-3-phosphocholine (POPC) bilayer using the Drude polarizable force field (FF). Our approach including both unbiased and umbrella sampling simulations. By using a polarizable FF, we sought to investigate explicit dipole responses in relation to local electric fields along the membrane normal. We evaluate molecular dipole moments, which exhibit variation based on their localization within the membrane, and compare the outcomes with analogous simulations using the nonpolarizable CHARMM36 FF. This comparative analysis aims to discern characteristic differences in the free energy surfaces of permeation for the various amino acid analogs. Our results provide the first systematic quantification of the impact of employing an explicitly polarizable FF in this context compared to the fixed-charge convention inherent to nonpolarizable FFs, which may not fully capture the influence of the membrane dielectric gradient.

## INTRODUCTION

Phospholipid membranes mediate the passive diffusion of molecules in and out of cells and serve as the solvent for membrane proteins that perform essential cellular functions. Membrane proteins are involved in many different biological processes, such as enabling signal transduction or receptor-mediated and active transport of materials in and out of the cell. Membranes and membrane proteins interact with endogenous signaling molecules such as neurotransmitters and hormones, as well as pharmaceuticals and other xenobiotics. In fact, around 60% of all FDA-approved medications have targets at the cell surface.^1^ As such, understanding the physicochemical properties and dynamics of membranes, and subsequently their interactions with membrane proteins, is of critical importance.

Many investigations of membranes and membrane proteins have made use of molecular dynamics (MD) simulations and span a wide variety of systems. The earliest atomistic studies investigated the basic structural characteristics of lipids and lipid organization, but were restricted by the timescales that where attainable with the hardware available at the time.^2,3^ As computational efficiency increased and as force fields (FFs) became more sophisticated, simulating more complex membrane systems became more tractable. The role of MD simulations in studying membrane proteins has several been reviewed elsewhere,^4,5^ but as examples, these advancements enabled work in coarse grained simulations of self-assembling systems,^6^ studies of small molecules in bilayers,^7^ and all-atom studies of membrane proteins in phospholipid bilayers.^8^ In the past 20 years, simulations of membrane proteins have progressed from the picosecond timescale to simulations of a G protein-coupled receptor (GPCR) on the scale of tens of microseconds.^9^ This dramatic increase in accessible timescale has enabled studies of the atomistic details of GPCR activation,^10^ ion-membrane protein allostery,^11^ and membrane protein-ligand binding pathways,^12^ among others.

Despite all of these advancements, a persistent challenge in modeling membranes is related to the intrinsic dielectric gradient that exists as a function of position along the membrane normal. To date, the vast majority of published MD studies of phospholipid membranes and the molecules that interact with them have applied nonpolarizable FFs, thereby only approximating electronic polarization effects. Polarizable FFs have been maturing over the course of the last 20 years, with FFs such as Drude classical oscillator^13–18^ and AMOEBA multipole-induced dipole^19–22^ being developed. An area of emphasis in studies and reviews of MD in membrane systems is how employing polarizable FFs may provide more accurate descriptions of these molecular systems, given their to the ability to better capture lipid bilayer electronic properties.^15,17,23–25^ When considering the early applications of polarizable FFs, the Drude FF was shown to be more accurate in representing membrane potential compared to the CHARMM FF, which was too positive in comparison.^14^

The development of the Drude lipid FF began with the parametrization of dipalmitoylphosphatidylcholine (DPPC) in 2013,^15^ followed by extension to additional zwitterionic lipid types in 2017,^18^ and a subsequent update in 2023.^26^ As a result, it has become possible to simulate multiple types of saturated and unsaturated zwitterionic lipids, such as 1-palmitoyl-2-oleoyl-*sn*-glycero-3-phosphocholine (POPC) and 1-palmitoyl-2-oleoyl-*sn*-glycero-3-phosphoethanolamine (POPE), with explicit electronic polarization. Recently, polarizable simulations of ion permeation^27^ and cation transport through the gramicidin A (gA) channel^28^ were reported. Work by Chen *et al.* on ion permeation mechanisms through bilayers of different thicknesses showed ion translocation consistent with experiments, and in better agreement than the results of nonpolarizable simulations.^27^ Similarly, Drude polarizable simulations of gA yielded better agreement with experimental conductance measurements than nonpolarizable simulations of gA.^28^ These studies show that explicit electronic polarizability is not only practical for membrane studies, but that the use of a polarizable model better captures the energetic and electronic properties of the membrane and proteins within it.

To further develop our understanding of the importance of electronic polarization effects in phospholipid membranes, we sought to characterize how using a polarizable FF impacts the interactions between membranes and small molecules derived from amino acids. Similar to prior studies,^29–33^ we studied amino-acid analogs and simulated them in a POPC bilayer using both the CHARMM36 (C36) nonpolarizable FF^34^ and the Drude polarizable FF^16,18^ to determine the role of electronic polarization on small-molecule localization, dipole moment changes, free energy surfaces, and electric fields acting within the membranes. Understanding the details of how these properties vary as a function of electronic polarization, using two FFs with similar target data and parametrization methodology, will help provide a more complete view of the underlying electrostatic forces acting within membrane systems.

## METHODS

### System Construction

The bilayer used for all simulations was generated using the CHARMM-GUI Membrane Builder.^35–40^ A total of 7200 TIP3P^41–43^ waters and 144 POPC lipids (72 in each leaflet) were generated. Coordinates for the amino-acid sidechain analogs were constructed using the CHARMM^44^ internal coordinate builder and then inserted into the center of the bilayer (Figure 1A). No ions were added to systems in which the small molecules carried no net charge. In cases where the small molecules were charged, a single counterion (1 K^+^ for acetate and 1 Cl^-^ for methylguanidinium) was included in the aqueous phase to neutralize the net charge of the system. We simulated a total of eight systems with different small molecules, including hydrophobic and aromatic (isobutane, benzene, and indole), polar neutral (methylimidazole, ethanol, and ethanethiol), and charged (acetate and methylguanidinium) species (Figure 1B).

**Figure 1.**
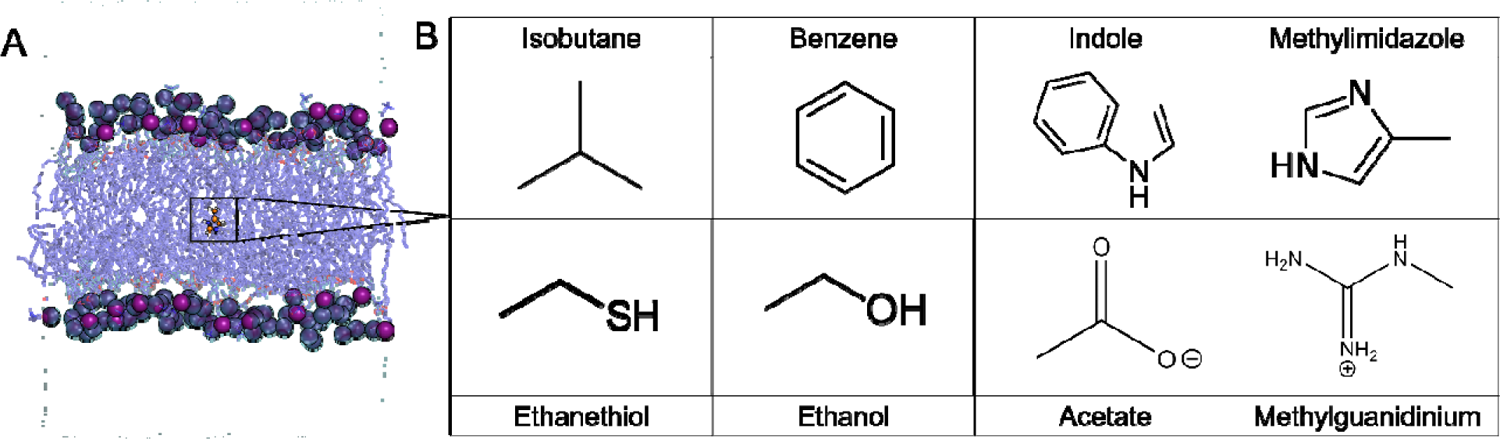
Schematic summary of systems. (A) Example of the initial configuration of each system. The small molecule inserted at the center of the bilayer is represented as orange ball-and-stick, lipids tails as dark blue sticks, phosphorus atoms as purple spheres, and water as cyan surface. (B) Structures and common names of each small-molecule sidechain analog.

### Unbiased MD Protocol

All simulation systems were initially prepared using the additive C36 FF. All systems were energy minimized using 1000 steps of steepest-decent and 1000 steps of adopted-basis Newton-Raphson (ABNR) minimization in CHARMM. Following minimization, equilibration was performed in NAMD for 1 ns,^45^ during which time position restraints were applied to the non-hydrogen atoms of the sidechain analogs with a force constant of 5 kcal mol^-1^ Å^-2^. Three independent simulations were initiated with different starting velocities under an NPT ensemble. The system temperature was regulated at 298 K with the Langevin thermostat method^46,47^ with a friction coefficient, γ, of 5 ps^-1^ and the pressure set to 1 atm with the Langevin piston method^46^ (oscillation period = 200 fs, decay time = 100 fs). Bonds involving hydrogen atoms were kept rigid with the SHAKE algorithm,^48^ allowing for a 2-fs time step in the C36 simulations. The short-range van der Waals forces were smoothly switched to zero from 10 to 12 Å, and periodic boundary conditions applied in all spatial directions. To calculate the electrostatic forces, the particle mesh Ewald^49,50^ (PME) method was used with a real-space cutoff of 12 Å and a grid spacing of approximately 1 Å.

We converted the coordinates of the equilibrated C36 systems to the Drude FF by adding Drude oscillators and lone pairs using the CHARMM program.^44^ Corresponding topologies were generated using the Drude-2019 protein^16^ (model compounds) and Drude-2017^18^ (lipids) parameter sets. The water molecules were converted from TIP3P to the polarizable SWM4-NDP^51^ water model. The Drude systems underwent an additional minimization to relax the positions of the Drude oscillators with other atoms held fixed. These systems were then equilibrated using NAMD for 1 ns as described above for the C36 simulations, except that the time step was set to 1 fs, the van der Waals potential was subjected to a switching potential, and a dual Langevin thermostat was used as described previously.^52,53^ Unrestrained simulations for all systems were performed using OpenMM.^54,55^ The duration of each replicate simulation was 1 μs, resulting in 3 μs of sampling per small molecule under each FF studied here.

### Umbrella Sampling

The equilibrated states for the unbiased simulations were used as the starting points for umbrella sampling. The reaction coordinate was defined as the z-axis component of the center-of-mass (COM) vector connecting the small molecule and the bilayer, denoted hereafter as Δz. For each system, we generated the initial configurations for each sampling window using CHARMM by translating the small molecule along the positive z-axis by 1 Å, resulting in 31 windows across the reaction coordinate, spanning values of Δz = 0 Å to Δz = 30 Å. Given the symmetry of the POPC membranes, we only generated half of the possible reaction coordinate describing passage of each small molecule across the membrane, as it should also be symmetric.

Each translation was followed by energy minimization and equilibration as described above for the unbiased MD simulations. These coordinates served as a starting point for equilibration using OpenMM, during which time position restraints were applied to the heavy atoms of the small molecules as described above. In this case, the Monte Carlo membrane barostat was used to employ semi-isotropic scaling, allowing for uniform scaling in the x-y plane while allowing the z-dimension to be scaled independently. This barostat was used in conjunction with the Langevin integrator using the same settings as described in the unbiased MD protocol. The restraint force constant was set to 1000 kJ mol^-1^ nm^-2^ to match the force constant used for subsequent umbrella sampling. The equilibrated C36 coordinates of each window were converted to the Drude convention, minimized, and equilibrated in OpenMM as described above.

The biasing potential applied in umbrella sampling was implemented using the CustomCentroidBondForce in OpenMM, applying a harmonic restraint force constant, *k*, of 1000 kJ mol^-1^ nm^-2^ between the z-axis component of the COM of the bilayer and the COM of the small molecule for most systems. The benzene system required a larger force constant of 1500 kJ mol^-1^ nm^-2^ for the Drude system to achieve adequate window overlap, as will be discussed below. Each window was simulated for at least 50 ns, and extended as needed in cases for which the free energy surfaces did not converge within this time. Convergence criteria are described below in the “Analysis” section. A complete listing of the simulation times in each window for each system is provided in the Supporting Information, Table S1. Convergence plots are also available in the Supporting Information, Figure S1 and S2 for C36 and Drude respectively.

### Analysis

The free energy surface for each system was generated with the weighted histogram analysis method (WHAM).^56,57^ To assess convergence, we computed the surfaces in 10-ns intervals over the last 40 ns of simulation time. If each of the 10-ns free energy surfaces were within the standard deviation of the free energy surface of the contiguous 40-ns time, the simulation was considered adequately converged. Otherwise, the simulations in each window were extended until convergence was achieved.

Electric field analysis was performed using the TUPÃ program.^58^ Electric field magnitudes were calculated in 10-ns intervals, each of which produced a mean value. Standard deviations were calculated from the mean values of each of these time intervals. The environment for the electric field calculation was defined as all lipids and any water molecule within 10 Å of the center-of-geometry (COG) of each small molecule through the ‘Include_Solvent’ functionality of TUPÃ. Dipole moment and localization analysis were performed in CHARMM.^44^ Molecular dipole moments were computed at the COM of each species, which eliminates the dependence on absolute coordinates in the case of charged species.

Hydration analysis was completed by using MDAnalysis^59,60^ to count the number of water oxygen atoms within 5 Å of the COM of each small molecule. Hydrogen bond analysis^61^ was also performed with MDAnalysis, using a distance cutoff of 3.5 Å and donor-hydrogen-acceptor angle cutoff of 150°.

## RESULTS AND DISCUSSION

### Aliphatic and Aromatic Small Molecules

Unbiased simulations of isobutane in POPC produced with both the C36 and Drude FFs resulted in similar localization profiles (Figure 2A). We define the position of the small molecule in terms of Δz, the z-component of its relative COM distance from that of the center of the membrane. Under this convention, Δz = 0 Å corresponds to the small molecule being exactly coincident with the center of the membrane along the z-axis. Isobutane largely remained within the bilayer core over the course of the simulations, as expected, though the distribution of Δz was slightly narrower using the Drude FF. In the polarizable simulations, the molecular dipole moment of isobutane varied considerably as a function of proximity to the membrane interface (Figure 2B). In the center of the bilayer, isobutane had an average dipole moment of ∼0.3 D, slightly polarized relative to its gas-phase value of 0.225 D,^62^ reflecting a small polarization response due to the low dielectric environment in the acyl chain region of the membrane. As isobutane sampled positions within the glycerol region, its dipole moment increased to >0.4 D, even sampling values in excess of 0.6 D (Figure 2B). In contrast, when simulated with the C36 FF, isobutane had a constant dipole moment on average, with only very small variations that can be attributed to bond vibrations and minor configurational fluctuations rather than a true change in electronic structure. Isobutane did not exit the bilayer during the Drude simulations, but did so briefly in the C36 simulations, thus resulting in sparse sampling within the headgroup region of the lipid bilayer (Figure 2A,B).

**Figure 2.**
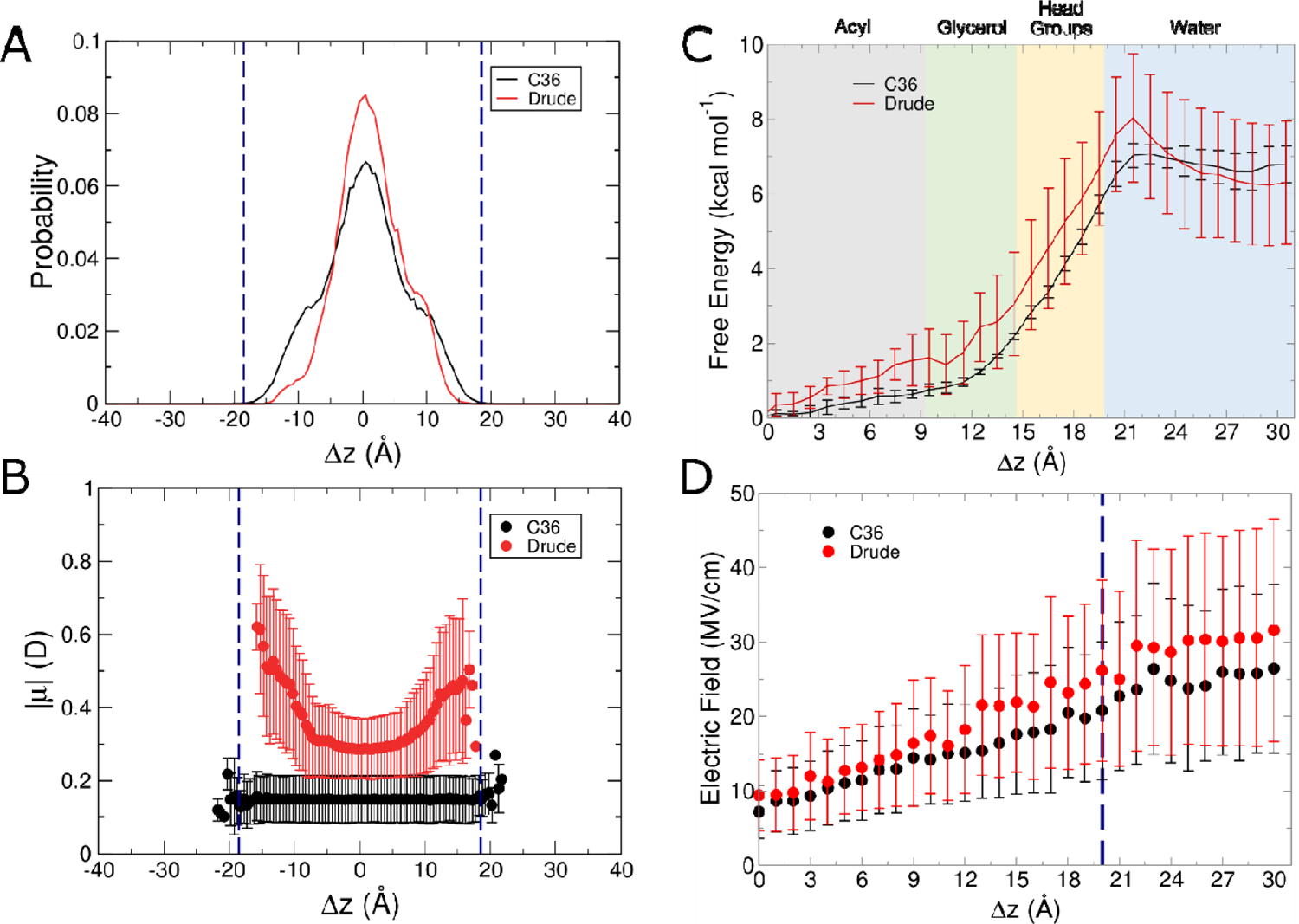
Results from C36 and Drude simulations of isobutane systems. (A) Normalized probability of localization of isobutane in unbiased simulations. (B) Molecular dipole moments as a function of position within the membrane in unbiased simulations. Error bars correspond to the root-mean-squared fluctuation (RMSF) of the binned data points. (C) Free energy surfaces from umbrella sampling simulations. (D) Average intrinsic electric field magnitude acting at the COG of isobutane in each umbrella sampling window. Blue dashed lines in panels (A), (B), and (D) denote the average position of the phosphate groups.

To more fully characterize the behavior of isobutane as it permeates into the POPC membrane and to more fully capture its properties in the aqueous phase, we performed umbrella sampling simulations along the membrane normal. The free energy surfaces produced from these simulations using both FFs are shown in Figure 2C. The global energy minimum in each surface is coincident with the center of the bilayer. The exit of isobutane from the membrane in characterized by a barrier of about 7 kcal mol^-1^ with both FFs, and the free energy plateaus throughout the aqueous layer. As isobutane moves along the membrane normal, it encounters increasing dielectric values, which can also be reflected in the electric field exerted on it. Figure 2D shows the average magnitude of the intrinsic electric field exerted by lipids and water on isobutane in each window of the umbrella sampling simulations. For both C36 and Drude, the average magnitude increases as isobutane transitions from the center of the bilayer to bulk water. Starting between the glycerol and head group regions, the field exerted in the Drude simulations is slightly larger than that of the C36 simulations, though both quantities are within error of one another, however the results suggest a polarization effect that is absent in the nonpolarizable simulations.

Given that isobutane is a nonpolar molecule with a small permanent dipole (gas phase μ = 0.225 D with the Drude FF), it is not surprising that the two FFs ultimately produced similar free energy surfaces. That is, the influence of electronic polarization in this case may be fairly small. Alkanes typically have relatively large molecular polarizability values, but this property only imparts a somewhat small change to the intrinsically small permanent dipole moment. Therefore, the properties of such species may in fact be well modeled by nonpolarizable FFs.

Interestingly, in the unbiased C36 simulations, isobutane diffused to the membrane-water interface and occupied this region for a few frames, suggesting that with the C36 FF, this molecule could interact with the aqueous environment more readily than with the Drude FF. It is possible that the absence of polarization in the lipids themselves leads to a somewhat weaker interaction between isobutane and the membrane, allowing for more rapid movement to the interface, though this location is ultimately disfavored compared to the bilayer core. Thus, isobutane appears to be a species for which the C36 and Drude FFs yield similar free energy surfaces and overall behavior. This result implies that either explicit electronic polarization is not crucial for modeling small, nonpolar compounds in membrane environments, or that there is serendipitous error cancellation in the C36 FF with respect to alkyl properties (the small molecule and the lipid bilayer). As noted in the original derivation of the Drude parameter set for alkanes, the CHARMM force field yields equivalent values for the dielectric constants of different alkanes, whereas the Drude FF better models the subtle differences among these chemical species.^62^ The better representation of dielectric constants should imply the importance of electronic polarization, even for these aliphatic species. In the context of a membrane, for which the alkyl groups of the lipids and the isobutane molecule itself lack polarization and may have similar dielectric properties, this combination may produce reasonably accurate results, though the physical underpinnings may be somewhat lacking.

The next small molecule we analyzed was benzene, which, while also nonpolar, has a π electron system that gives rise to characteristic properties. As with isobutane, both the C36 and Drude FFs led to similar localization trends (Figure 3A) in the unbiased simulations. Benzene was largely found at the center of the bilayer or in the glycerol region; with C36, the distribution is more uniform than with the Drude FF, which has more distinct subpopulations at these locations. As with isobutane, using C36 led to sparse sampling of benzene outside of the bilayer, though the entire aqueous layer was sampled. In contrast to isobutane, in one replicate of the Drude simulations, the benzene molecule briefly sampled the space ∼1 Å outside of the bilayer. Also following the trend of isobutane, the dipole moment of benzene using the Drude FF was strongly dependent on where it was located in the bilayer, as shown in Figure 3B. The dipole moment of benzene was nearly twice as large when occupying the headgroup region and positions outside the bilayer compared to when it resided in the center of the bilayer. This behavior is similar to that of what was seen for isobutane, with the C36 FF resulting in a nonpolar species sampling locations outside of the bilayer. Paired with the isobutane results, this outcome shows a trend for hydrophobic species, wherein C36 producing sampling of what should presumably be unfavorable, high free energy states.

**Figure 3.**
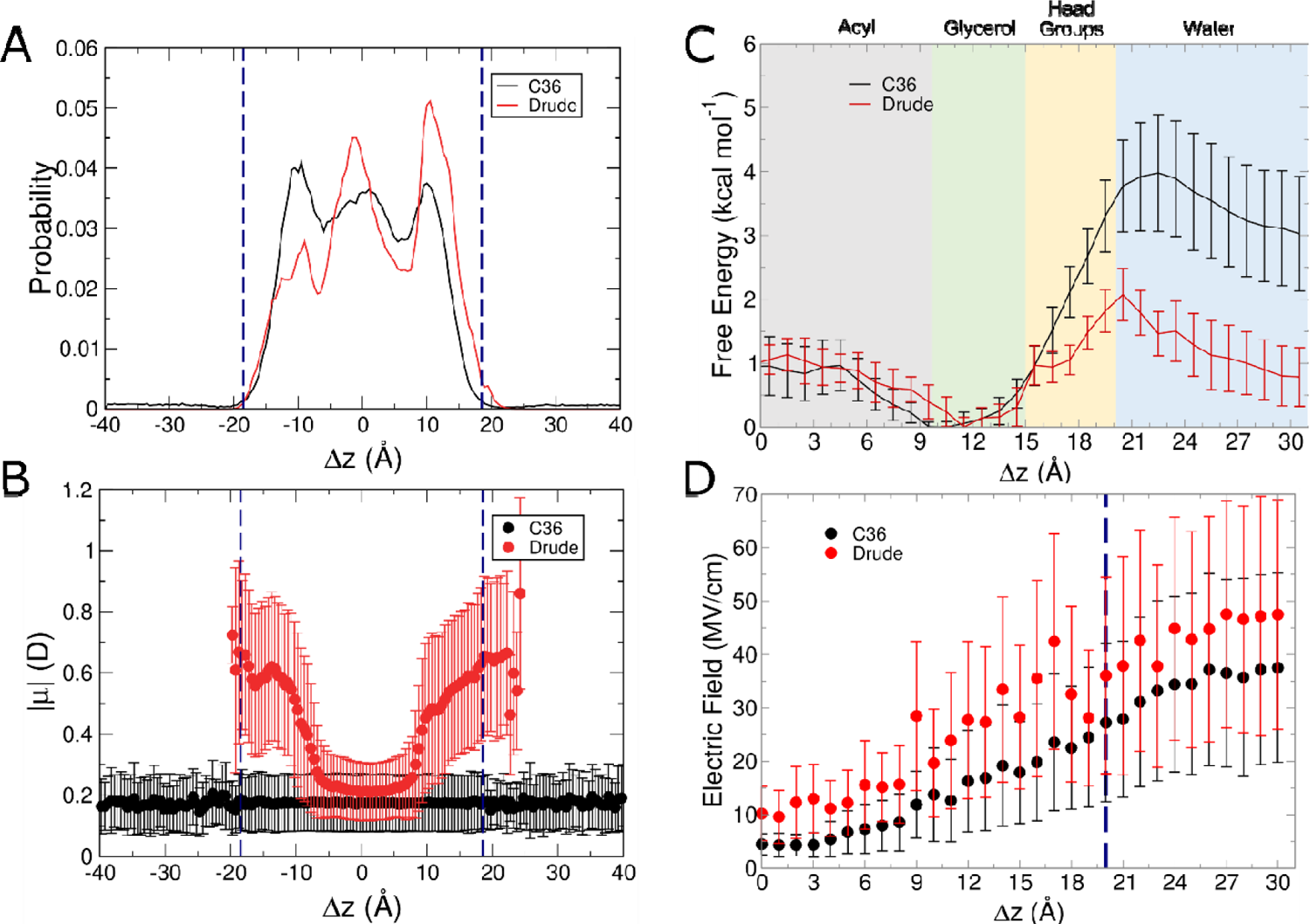
Results from C36 and Drude simulations of benzene systems. (A) Normalized probability of localization of benzene in unbiased simulations. (B) Molecular dipole moments as a function of position within the membrane in unbiased simulations. Error bars correspond to the RMSF of the binned data points. (C) Free energy surfaces from umbrella sampling simulations. (D) Average intrinsic electric field magnitude acting at the COG of benzene in each umbrella sampling window. Blue dashed lines in panels (A), (B), and (D) denote the average position of the phosphate groups.

The results of the umbrella sampling simulations confirm that the relative free energy of benzene in the aqueous layer outside the POPC bilayer is somewhat higher with C36 than for the Drude FF (Figure 3C). As with isobutane, the ability of benzene to access these higher-energy states may be due somewhat to the lack of polarization among the lipids and therefore faster diffusion of the benzene molecule into the aqueous layer, where it rapidly diffuses as it is unable to form favorable hydrogen bonding interactions with water. With the Drude FF, the relative free energy of occupying the aqueous layer is only ∼1 kcal mol^-1^ higher than the energy minimum within the glycerol region, suggesting that the explicit polarization of the benzene molecule allows for somewhat more favorable interactions with the polar solvent than is possible with C36.

In contrast to isobutane, which experienced similar electric fields with each FF until reaching the glycerol region, the electric field felt by benzene differed between the two FFs even at the core of the bilayer (Figure 3D). This finding suggests that even though both species, bearing small dipole moments with the C36 FF, felt slightly different electric fields from the lipids and water. Additionally, in the C36 simulations, benzene felt fields that ranged from ∼5 MV/cm in the core of the bilayer to ∼40 MV/cm in water, a much greater range than that of isobutane (∼10 MV/cm to ∼25 MV/cm), likely as a consequence of the greater magnitudes of partial charge on the atoms of benzene. This difference was even more distinct with the Drude FF. Isobutane and benzene both experienced electric fields of ∼10 MV/cm in the bilayer core, suggesting their electrostatic properties are comparable in this environment, but whereas isobutane was subjected to fields of ∼30 MV/cm in water, the field acting on benzene in water was much larger, ∼50 MV/cm. Thus, the influence of electronic polarization in these species becomes more distinct upon partitioning into the aqueous phase, and the π system of benzene likely contributes to a greater mutual polarization effect with water molecules.

Despite being generally considered nonpolar, the impact of electronic polarization in benzene is clearly much greater than in the case of isobutane. The electronic structure of benzene, with its delocalized π-electron cloud, leads to a concentration of negative charge on the faces of the ring. This property likely confers more specific, directional interactions with polar species like water and lipid headgroups that respond more strongly to induced polarization and therefore are important to model accurately in membrane systems.

The final molecule in the nonpolar and aliphatic group that we considered was indole, which in addition to being aromatic like benzene, is the first molecule analyzed that contains a heteroatom. As with the previous molecules, both FFs produced similar localization trends (Figure 4A). With both the C36 and Drude FFs, indole was enriched at the membrane-water interfaces, with only sparse localization in other regions. In the simulations with the Drude FF, however, indole had a slightly higher occupancy outside of the bilayer compared to the C36 simulations. We note that in the unbiased simulations, indole had an asymmetric distribution in the membrane, apparently favoring one side of the bilayer compared to the other. Over the course of the simulations, indole moved toward one bilayer leaflet, and then tended to stay there for the remainder of the simulation. Given that the bilayer we simulated was homogenous, with the same number of POPC per leaflet, we anticipated a more symmetric distribution and therefore suspected that the initial orientation of indole was biasing the outcome. We performed three additional replicates using an alternate orientation for indole, such that it was rotated 180° relative to the starting configuration of the previous simulations. The rotation oriented the NH group toward the other membrane leaflet. The results of these simulations are shown in Supporting Information, Figure S3, and represent improved symmetry in the distribution of indole along the *z*-axis, suggesting that the apparent asymmetry observed in the original simulations does not correspond to a true preference but is a minor artifact due to the starting orientation of the molecule.

**Figure 4.**
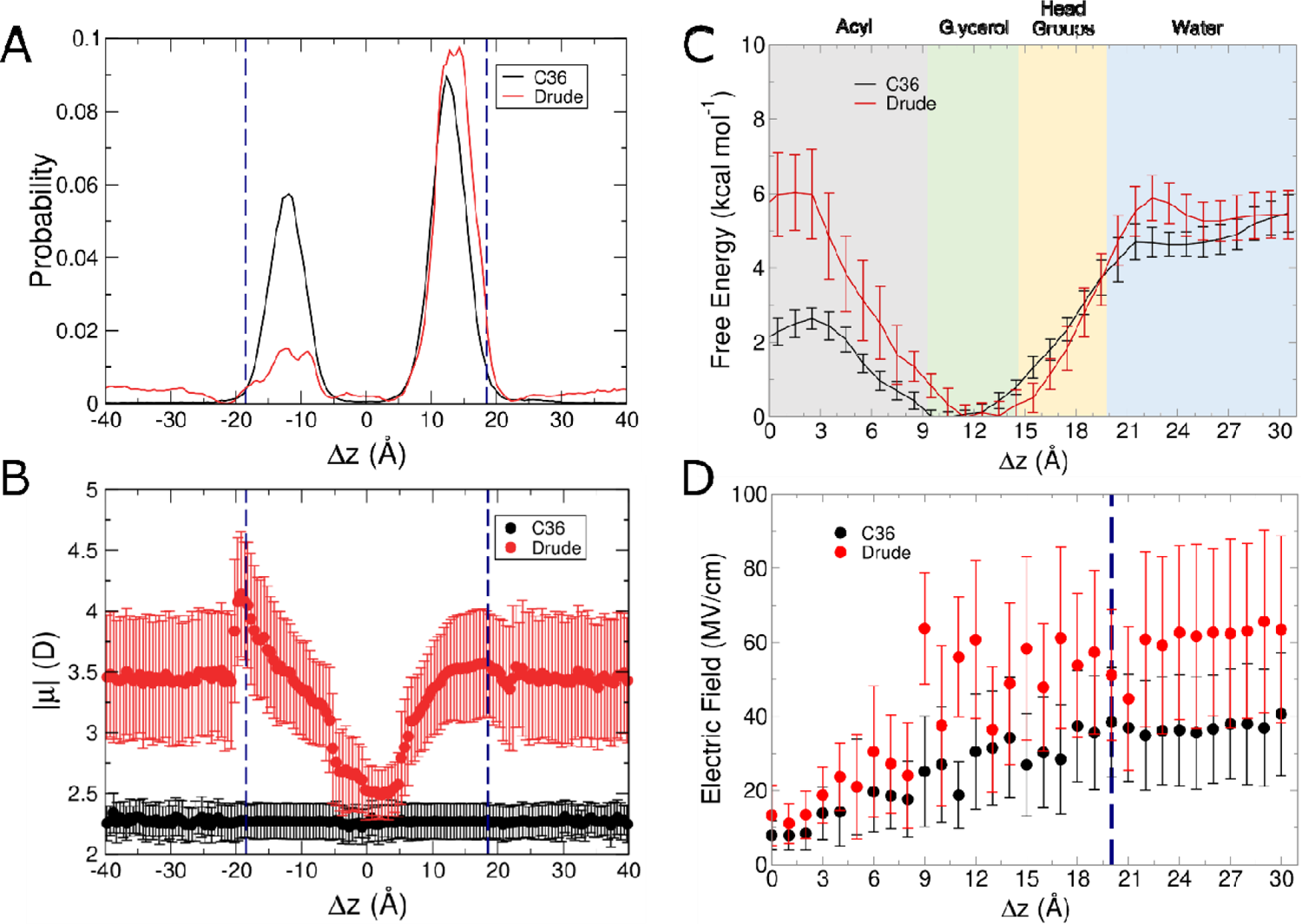
Results from C36 and Drude simulations of indole systems. (A) Normalized probability of localization of indole in unbiased simulations. (B) Molecular dipole moments as a function of position within the membrane in unbiased simulations. Error bars correspond to the RMSF of the binned data points. (C) Free energy surfaces from umbrella sampling simulations. (D) Average intrinsic electric field magnitude acting at the COG of indole in each umbrella sampling window. Blue dashed lines in panels (A), (B), and (D) denote the average position of the phosphate groups.

As with the previous molecules, the dipole moment of indole was dependent on its location in the bilayer when using the Drude FF but not with the C36 FF, as shown in Figure 4B. In the glycerol regions of the bilayer, the dipole moment of indole (∼3.5 D) was greater by 1 D compared to when it was located at the center of the bilayer (∼2.5 D). Values as high as 4 D were sampled in the headgroup region. This value is comparable to the permanent dipole moment of gas-phase indole in a monohydrate complex, obtained via spectroscopic measurements.^63^ With the C36 FF, the dipole moment of indole was similar to the Drude gas-phase dipole moment of 2.25 D and MP2/6-31G* calculations of 2.21 D.^64^ As such, indole is modeled as being somewhat more hydrophobic than with the Drude model, resulting in less favorable interactions with water, thus the difference in sampling seen between Drude and C36 with the C36 simulations yielding fewer instances of direct interaction with water.

In the umbrella sampling simulations, the Drude FF produced a higher barrier to being directly in the center of the bilayer compared to C36, and ultimately a similar barrier to exit the bilayer (Figure 4C). The minimum is shifted slightly deeper into the bilayer for C36, whereas with the Drude FF, it coincides with the center of the glycerol region. Interestingly, the locations of the free energy minima for both benzene and indole were similar with each FF. That is, for C36, the minima for benzene and indole were both around Δz = 9 Å, but with the Drude FF, the respective minima were around Δz = 11 Å. The minima differ, however, in their shapes. In the case of benzene, the Drude FF yielded very small barriers for deeper permeation of benzene into the bilayer and entry into water (∼1 and ∼2 kcal mol^-1^, respectively). In contrast, indole had very steep barriers to either side, suggesting that while the localization at the glycerol interface was favorable for both of these aromatic molecules, indole is more strongly driven to this location. This distinction was not present with the C36 FF, for which the free energy barrier to deeper permeation were on the order of 1-2 kcal mol^-1^ and for entry into water, ∼4 kcal mol^-1^ for both species.

Considering the aromatic nature of indole, its localization and corresponding free energy surface with both FFs are generally what is expected for aromatic compounds, particularly in the context of membrane proteins, which contain a characteristic “aromatic belt” that features in insertion, embedding, and stabilization in the bilayer.^65^ When considering the small molecules and where their analogs are generally found, tryptophan is generally present in protein structures such that they are enriched at interfacial regions, whereas phenylalanine is more associated with the hydrophobic core of membrane proteins.^66^ This preference is reflected in the localizations and free energy surfaces of their corresponding analogs with the Drude FF, given that indole is more strongly driven into the glycerol interface region with a much larger free energy barrier to further permeation into the acyl chain region for indole. Indole has a larger molecular polarizability than benzene (12.93 Å^3^ and 8.3 Å^3^ with the Drude FF, respectively), which could explain this difference, particularly in the context of the electric fields that the molecules experience in the different membrane microenvironments. In the simulations of indole, the intrinsic electric fields exerted on it follow the same general trend as benzene, starting around 10 MV/cm in the bilayer core, and increasing to above 50-60 MV/cm around the glycerol interface and 60 MV/cm in water. The latter two values are much higher magnitude than the fields acting on benzene. This outcome is likely due to the larger molecular polarizability of indole (reflecting greater sensitivity to electric fields and thus a larger mutual polarization effect on the lipids and water), its capacity to form hydrogen bonds via its pyrrole ring, and potential for cation-π interactions.^67^ We counted the average hydrogen bond count occurring between the water and indole, and ultimately there was largely similar hydrogen bond counts occurring when comparing C36 and Drude (Supporting Information, Figure S4A).

### Polar, Neutral Small Molecules

The first of the polar, neutral small molecules we analyzed was methylimidazole. Similar to indole, methylimidazole had a higher probability of occupying the interfaces with the Drude FF compared to simulations performed with C36 (Figure 5A). With the C36 FF, methylimidazole was locally enriched at the interface but was more likely to be in the aqueous phase (Figure 5A). The dipole moment continued to follow the trend of being larger at the interfaces, and smaller at the interior of the bilayer for Drude (Figure 5B). Among the molecules studied here, methylimidazole is the first for which the Drude FF produced a lower dipole moment than C36 at the center of the bilayer. The only other instance of such a phenomenon is in the case of ethanol (discussed below). Further, methylimidazole had the largest change in dipole moment of all simulated molecules, increasing by ∼2 D as it moved from the center of the bilayer into water. Given similar behavior to indole in terms of localization asymmetry, we opted to simulate additional replicates with methylimidazole flipped 180° (Supporting Information, Figure S5). This orientation led to somewhat more symmetric sampling in the Drude simulations, suggesting that the apparent asymmetry observed in the original simulations does not correspond to a true preference but is a minor artifact due to the starting orientation and sampling.

**Figure 5.**
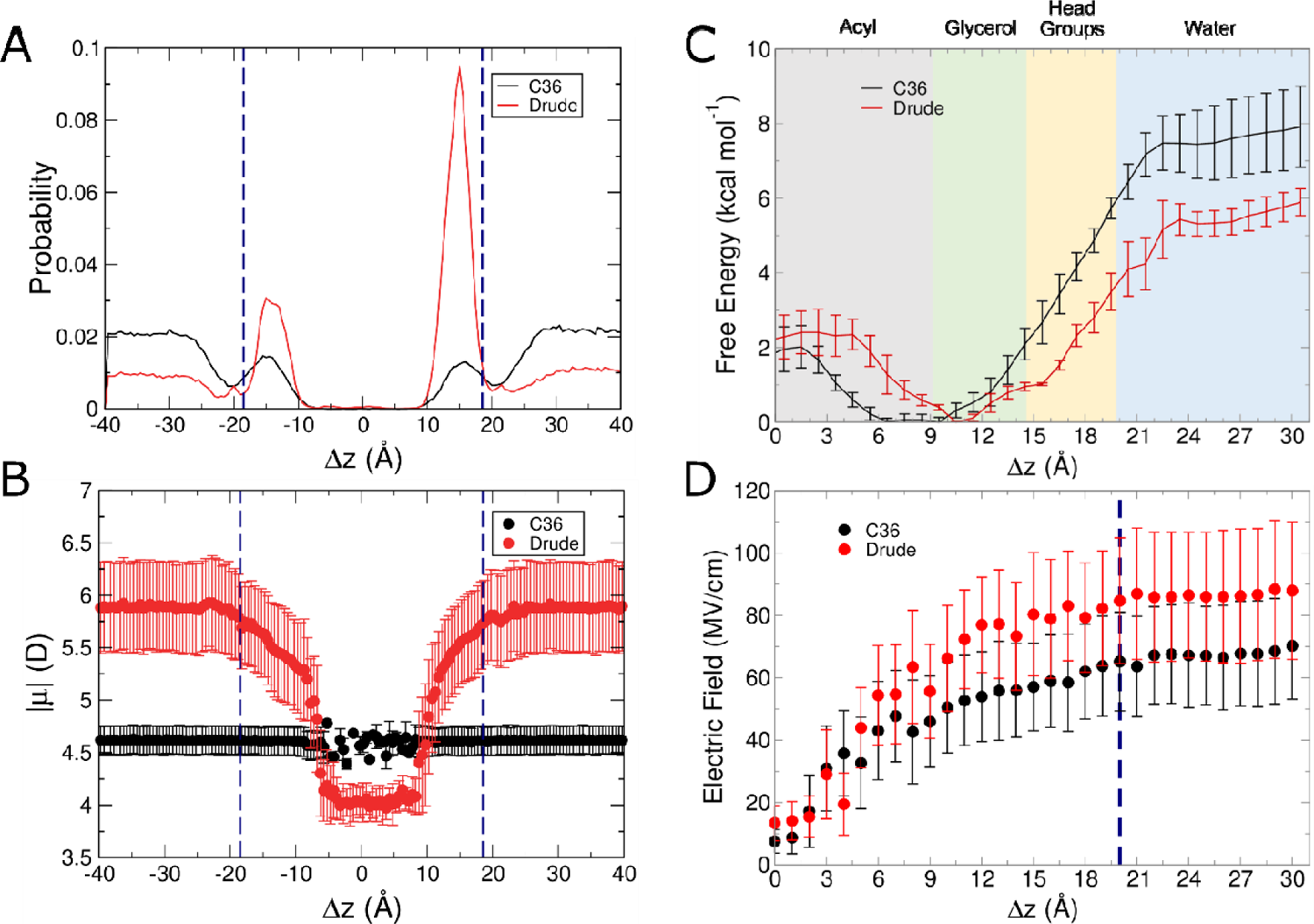
Results from C36 and Drude simulations of methylimidazole systems. (A) Normalized probability of localization of methylimidazole in unbiased simulations. (B) Molecular dipole moments as a function of position within the membrane in unbiased simulations. Error bars correspond to the RMSF of the binned data points. (C) Free energy surfaces from umbrella sampling simulations. (D) Average intrinsic electric field magnitude acting at the COG of methylimidazole in each umbrella sampling window. Blue dashed lines in panels (A), (B), and (D) denote the average position of the phosphate groups.

Methylimidazole is also aromatic, and therefore behaved similarly to indole and benzene, such that its free energy minimum was located at the glycerol interface with the Drude FF but deeper within the bilayer with the C36 FF (Figure 5C). The magnitude of the intrinsic electric field acting on methylimidazole was 10 MV/cm at the center of the bilayer (similar to both benzene and indole), and increased out into water to around 90 MV/cm for the Drude FF, with the C36 FF exerting a weaker field (Figure 5D). This outcome builds upon the observation in the case of indole, in that methylimidazole has the ability to form two hydrogen bonds instead of one, thus it is likely that this molecule more strongly induces dipole responses in nearby molecules, including the glycerol moieties of POPC and water. When considering the average count of hydrogen bonds per window, ultimately there is a similar amount when comparing C36 and Drude (Supporting Information, Figure S4B). However, we do see a higher average count when comparing methylimidazole to indole, underlining the difference in their ability to hydrogen bond.

The final two polar uncharged molecules we analyzed were ethanethiol and ethanol. Ethanethiol served as a small, polar, sulfur-containing species, with ethanol as its alcohol analog. As these two species are the two most similar molecules we studied, we discuss them together. Figure 6A shows the unbiased localization of ethanethiol, which was similar to that of benzene, exhibiting a broad distribution with small peaks at the center of the bilayer and each of the two glycerol regions. The highest probability of occupancy with the Drude FF occurred at the interfaces, whereas the C36 FF resulted in a slight preference for the center of the bilayer, though its peaks were less distinct than those produced with the Drude FF. Figure 6B shows the dipole moment, which follows the trends of all prior small molecules, with the polarizable model of ethanethiol increasing its dipole moment by ∼0.5 D from the center of the bilayer to the interface. The free energy surfaces follow the same trend with both FFs. The minimum is coincident with the center of the bilayer, but interesting differences emerge as the small molecule approaches the aqueous phase. Overall, the Drude FF predicts a higher barrier to membrane exit; in other words, ethanethiol is more strongly driven into the bilayer with the polarizable FF (Figure 6C). With C36, the free energy surface exhibits a broad, shallow local minimum in the glycerol region, a feature that is absent from the Drude surface.

**Figure 6.**
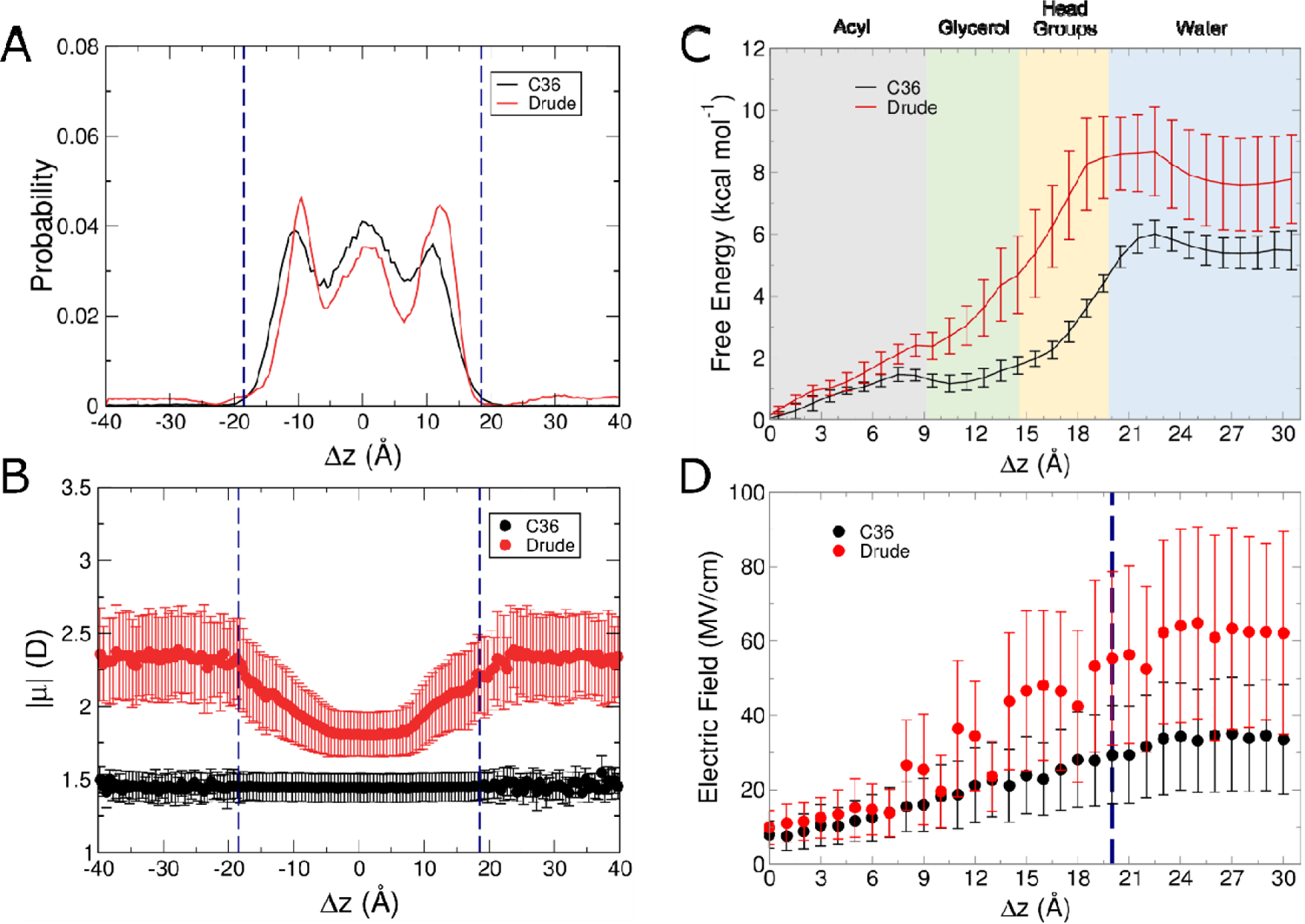
Results from C36 and Drude simulations of ethanethiol systems. (A) Normalized probability of localization of ethanethiol in unbiased simulations. (B) Molecular dipole moments as a function of position within the membrane in unbiased simulations. Error bars correspond to the RMSF of the binned data points. (C) Free energy surfaces from umbrella sampling simulations. (D) Average intrinsic electric field magnitude acting at the COG of ethanethiol in each umbrella sampling window. Blue dashed lines in panels (A), (B), and (D) denote the average position of the phosphate groups.

The presence of this shallow local minimum in the glycerol environment and the overall lower barrier to membrane exit with the C36 FF suggests that the additive model of ethanethiol has a weaker driving force into the bilayer, and a flatter free energy surface than the that of the polarizable FF. However, in terms of the electrostatics, thiols are known to form weak hydrogen bonds, and as such are more hydrophobic than their alcohol analogs.^69^ Our simulations showed generally fewer than one hydrogen bond between ethanethiol and water, on average, in each umbrella sampling window (Supporting Information, Figure S4C). Whereas the C36 FF yielded essentially no hydrogen bonds, the Drude FF produced a slightly larger average, suggesting that the inclusion of lone pairs and explicit polarization (which is anisotropic in the case of the thiol sulfur atom) leads to a small increase in hydrogen-bonding capability of ethanethiol. Hydrogen bonding between the ethanethiol and POPC also differs between the force fields, with the Drude FF producing a few hydrogen bonds between ethanethiol and the ester groups, while C36 had none (Supporting Information, Figure S6A). The sum total of these interactions explains the energetic barrier to leaving the bilayer, as it may be more costly to disrupt these hydrogen bonds in the polarizable FF compared to the nonpolarizable FF.

The electric fields acting on ethanethiol are plotted in Figure 6D. Following the same trends of previous small molecules, the electric the fields acting on ethanethiol in the polarizable system were larger than with the C36 FF. Interestingly, the field magnitudes in the ethanethiol system were similar to those in the indole system (Figure 4D), suggesting that despite being chemically distinct and very different in size, the presence of a single hydrogen-bonding group within an otherwise aliphatic molecule may induce similar electric fields within the surrounding microenvironment.

Figure 7A shows the probability of finding ethanol at a given point in the bilayer. The localization with both FFs were similar, with local enrichment at the membrane-water interface, and considerable sampling within the aqueous phase. Molecular dipole moment behavior followed the trend of all other small molecules, with a smaller dipole moment at the center of the bilayer, and a larger dipole moment at the interfaces and exterior of the bilayer (Figure 7B). Similar to ethanethiol, ethanol is another case for which the free energy profiles differ prominently depending on the FF used (Figure 7C). With the Drude FF, the minimum is in the glycerol region, reflecting the preferred localization in the unbiased simulations. This localization is not surprising as neat ethanol and the glycerol region of phospholipid membranes have similar dielectric constants.^69^ Therefore, the glycerol interface environment may be optimal for accommodating ethanol, which has a short alkyl chain but can engage in hydrogen bonding via its hydroxyl group. In contrast, simulations with the C36 FF resulted in a free energy minimum at the very center of the bilayer, and subsequently a much higher barrier to membrane to exit than in the case of the Drude FF. This outcome seemingly contradicts the unbiased localization but can be explained by a greater degree of hydration occurring in the C36 simulations compared to the Drude simulations (Supporting Information, Figure S7C). With the C36 FF, ethanol became hydrated as deeply in the membrane as Δz = 2 Å, with more than 1 water hydrating ethanol on average occurring at Δz = 5 Å. Thus, the apparent discrepancy in terms of unbiased ethanol localization and free energy is explained by hydration that perturbs the free energy surface. With the Drude FF, on the other hand, there were essentially no water molecules within 5 Å of the COM of ethanol until Δz = 9 Å. By Δz = 11 Å, both FFs largely behaved similarly, but this outcome demonstrates a fundamental difference in hydration behavior in the bilayer for a polar molecule like ethanol.

**Figure 7.**
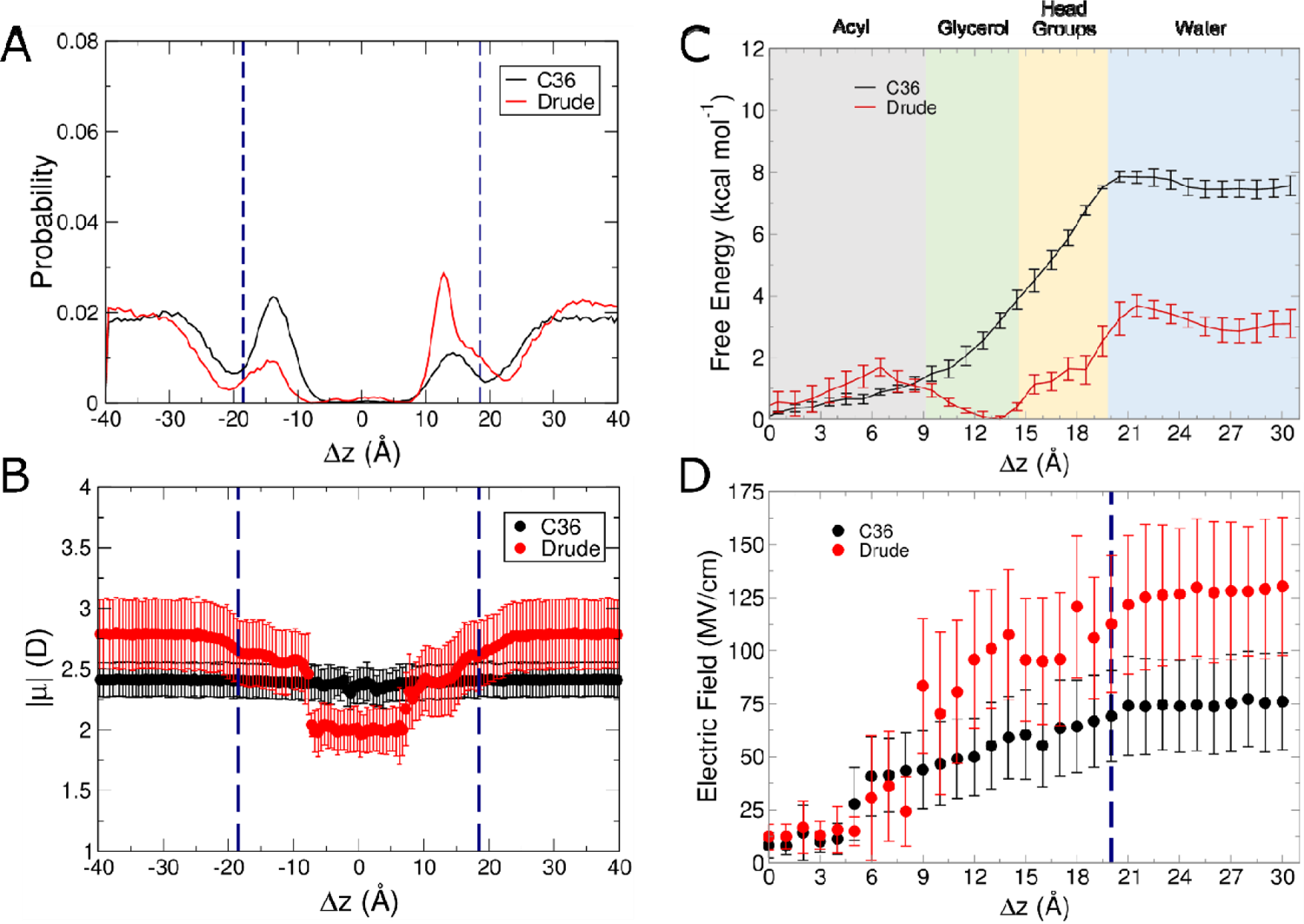
Results from C36 and Drude simulations of ethanol systems. (A) Normalized probability of localization of ethanol in unbiased simulations. (B) Molecular dipole moments as a function of position within the membrane i unbiased simulations. Error bars correspond to the RMSF of the binned data points. (C) Free energy surfaces from umbrella sampling simulations. (D) Average intrinsic electric field magnitude acting at the COG of ethanol in each umbrella sampling window. Blue dashed lines in panels (A), (B), and (D) denote the average position of the phosphate groups.

The hydration of ethanol with the Drude FF corresponds to the sharp increase in the electric field magnitudes (Figure 7D), as well as dipole moments (Figure 7B). The electric fields acting on ethanol were the highest among all the molecules considered here, reaching ∼125 MV/cm around the interface in the Drude simulations. Hydration of ethanol at this location likely contributes to the interfacial energy minimum with the Drude FF (Figure 7C). Given that it is a short-chain alcohol, ethanol is known to localize at the membrane-water interface rather than diffuse more deeply into the hydrophobic core.^70^ Hydrogen bonding with both water and the hydrophilic portion of the phospholipids likely govern this localization preference.

The degree of hydration and hydrogen bonding capabilities helps to explain subtle differences in the properties of ethanol with respect to its sulfur-containing analog. Ethanethiol is more hydrophobic and therefore engages in weaker hydrogen bonding than its alcohol counterpart. This phenomenon is apparent in both force fields studied, with hydration only occurring for ethanethiol in windows further from the bilayer center (Supporting Information, Figure S7B,C). Ethanethiol was effectively dehydrated until Δz = 11 Å with the Drude FF, while ethanol has clear hydration at Δz = 9 Å.

The free energy surfaces for ethanol and ethanethiol with the C36 FF were ultimately similar, though ethanol lacks the shallow local minimum at the interface that ethanethiol has. The greater degree of hydration of ethanol allows it to be accommodated more deeply in the bilayer with the C36 FF. Contrasting the nonpolarizable FF, ethanol localized at the interfacial region of the bilayer with the Drude FF, while ethanethiol was pushed more deeply into the bilayer, likely being driven by the difference in hydration and hydrogen bonding of the two species (Supporting Information, Figures S4C,D and S6A,B).

### Charged Small Molecules

Similar to ethanol, acetate was a case for which the free energy surface was very different depending on the explicit representation of polarization. As shown in Figure 8, acetate occupancy in the bilayer center is strongly disfavored with the Drude FF, with a barrier to permeation of 14 kcal mol^-1^. This outcome agrees with other umbrella sampling simulations of acetate in bilayers.^7,30,71^ In contrast, with the C36 FF, this trend is inverted, implying that acetate favorably resides in the bilayer core. This unexpected phenomenon was explained by aberrant hydration behavior. Over the course of the simulations in each umbrella sampling window, considerable water influx into the bilayer occurred in the case of C36. Some water penetrated the hydrophobic core of the membrane in the Drude simulations, but to a much smaller extent. The hydration perturbed the membrane structure (Figure 9) and complicates the interpretation of the free energy surface, because the POPC bilayer itself is in a very different state than in the other simulations performed here. The sparse hydration in the case of the Drude simulations indicates that acetate requires fewer hydrating water molecules to transit the membrane, likely as a response to the strong depolarization of the molecule in this location (Figure 8B). Thus, hydration behavior in ethanethiol, ethanol, and acetate systems are important for understanding the free energies surfaces and differences that arise as a function of including explicit electronic polarization.

**Figure 8.**
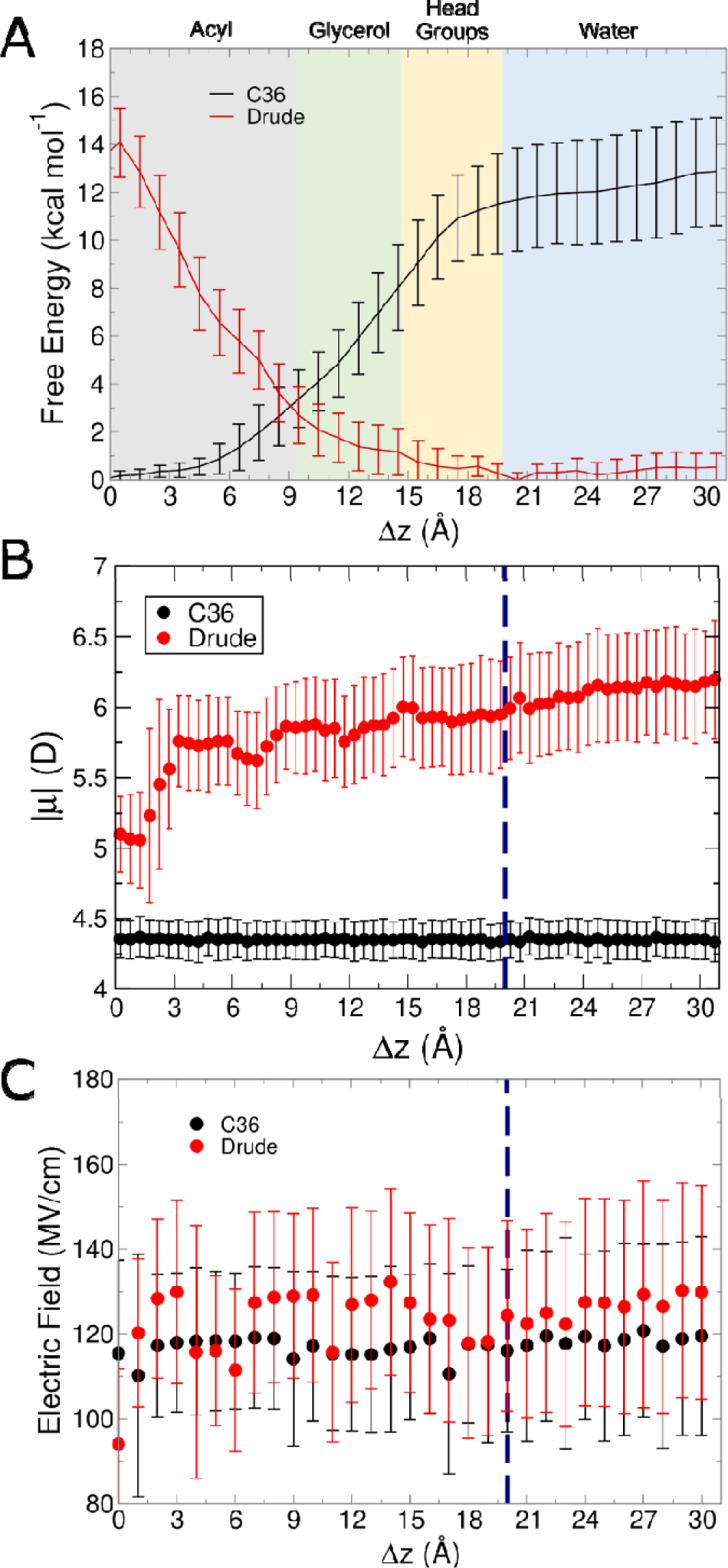
Results from C36 and Drude simulations of acetate systems. (A) Free energy surfaces from umbrella sampling simulations. (B) Molecular dipole moments as a function of position within the membrane in unbiases simulations. Error bars correspond to the RMSF of the binned data points. (C) Average intrinsic electric field magnitude acting at the COG of acetate in each umbrella sampling window. Blue dashed lines in panels (B) and (C) denote the average position of the phosphate groups.

**Figure 9.**
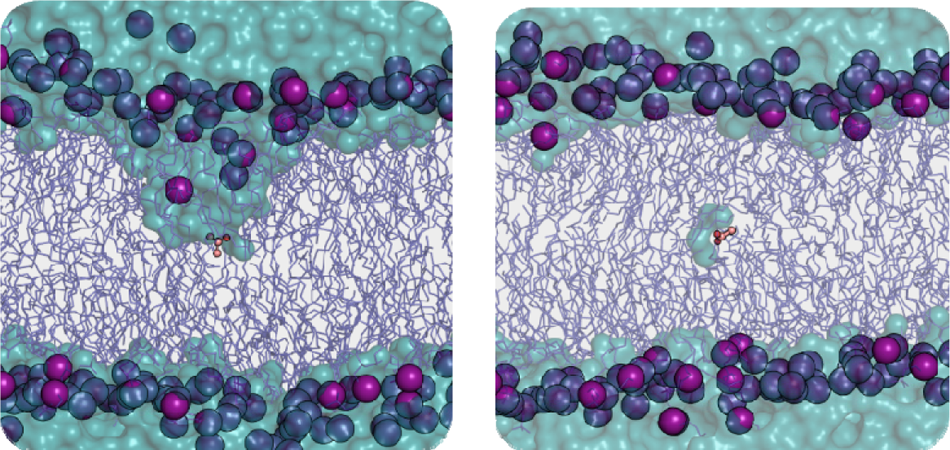
Hydration of acetate at Δz = 0 Å for the C36 (left, at 33 ns) and Drude FFs (right, at 38 ns). The acetate molecule inserted at the center of the bilayer is represented as orange ball-and-stick, lipids tails as dark blue sticks, phosphorus atoms as purple spheres, and water as cyan surface.

**Figure 10.**
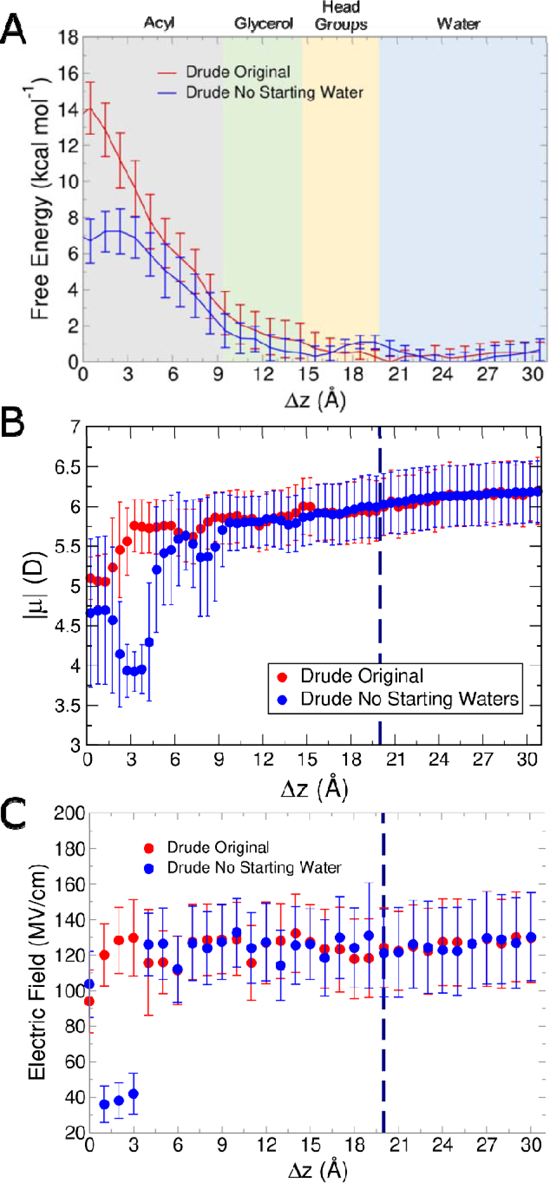
Acetate umbrella sampling results started from dehydrated coordinates compared to original starting coordinates (Drude only, Drude original repeated from Figure 9). (A) Free energy surfaces from umbrella sampline simulations. (B) Molecular dipole moments as a function of position within the membrane in unbiased simulations. Error bars correspond to the RMSF of the binned data points. (C) Average intrinsic electric field magnitude acting at the COG of acetate in each umbrella sampling window. Blue dashed lines in panels (B) and (C) denote the average position of the phosphate groups. Simulations ran for 70 ns to mirror original coordinates and achieved convergence with same applied methodology.

Figure 8C shows the intrinsic electric field acting on acetate, and we see the first difference in electric field trend among the small molecules. The electric field magnitudes felt by acetate were largely consistent across the windows, given that water influx occurred with both FFs. The lowest magnitude is at Δz = 0 Å for both FFs as this window had the lowest degree of hydration for both FFs (Supporting Information, Figure S7D). The average number of water molecules within 5 Å of the COM of acetate for the Drude FF was lower than the C36 FF at all values of Δz inside of the bilayer. The presence of this water provides an explanation for the difference in trends of electric fields, as well as potentially beginning to explain the difference in free energy surfaces between the two FFs.

Given that the general workflow of running a Drude simulation uses the final snapshot of C36 equilibration as the input coordinates for conversion to the Drude model, the presence of water in these states may have been an artifact of the additive FF. To test this possibility, we first performed an extended C36 equilibration with position restraints applied to acetate to ensure the same water influx occurred without the OpenMM CustomCentroidBondForce applied. The same membrane distortion and water influx were observed, making it unlikely that the external restraint force destabilized the membrane structure to allow water diffusion into the hydrophobic core.

We then converted the pre-equilibrated C36 coordinates to the Drude convention and equilibrated those as described in the Methods, starting the production simulation without any water within the bilayer, thus removing any bias of potential artifacts from the additive FF. We found that similar hydration occurred for most umbrella sampling windows, except for Δz = 1 – 3 Å (Supporting Information, Figure S7D). In those cases, essentially no water molecules were present within 5 Å of the acetate molecule, resulting in a lower barrier to entry of the bilayer, a lower dipole moment, and weaker electric field acting on the small molecule (Figure 10). Thus, acetate permeation is uniformly disfavored with the Drude FF, as expected, although the barrier is somewhat lower without any water associated with the acetate molecule.

Many studies have sought to understand ion translocation in bilayers, both through simulations^27,72,73^ and experiments,^74,75^ and there are thought to be two potential mechanisms, depending on bilayer thickness. The ion-induced mechanism occurs when the ion permeating the bilayer results in substantial membrane defects with water entering the bilayer core, while the solubility-diffusion mechanism results in the molecule first permeating into the hydrophobic core, then diffusing out to the other side with no membrane defect formation.^75^ It is thought that in thinner bilayers, the ion-induced defect mechanism is dominant, but as bilayer thickness increases, there is a transition to the solubility diffusion mechanism.^74^ In this case, the initial Drude simulations support the solubility diffusion mechanism, but the results with the C36 FF indicate the ion-induced defect mechanism. This outcome follows previous work comparing C36 and Drude,^27^ as well as experimental conclusions,^74,75^ which suggest that charged molecules transition from ion-induced defect to solubility diffusion at lipid chain lengths similar to POPC, which was used here. A definitive conclusion regarding ion permeation mechanisms from MD simulations would likely require more complex methods like transition-tempered metadynamics.^76,77^

It should also be noted that acetate would likely be protonated inside the bilayer. In the case of its amino-acid analog, aspartate, the protonation state can help drive membrane protein positioning, transmembrane region insertion, and regions that may be bound to the surface.^78^ Here, our goal was to investigate what barriers exist for charges permeating into bilayers when using a polarizable FF compared to a nonpolarizable FF. Doing so sets the basis for future investigations into the effects of protonation, which may require a combination of advanced techniques, such as constant-pH simulations,^79^ which are not yet compatible with the Drude FF.

The final small molecule we investigated was methylguanidinium, which is positively charged at neutral pH. As shown in Figure 11A, methylguanidinium is a case for which explicit polarization did not impact the free energy surface. The free energy maxima are located at the center of the bilayer with both FFs and their minima are in the glycerol/headgroup transition, with a gradual slope driving methylguanidinium into the respective minima. Figure 11B show the dipole moment change, where C36 simulations remained flat, and methylguanidinium slightly depolarized upon exiting the bilayer in the Drude simulations. The electric fields acting on methylguanidinium also are flat (Figure 11C), indicating the electronic environment is largely consistent across the umbrella sampling windows. This phenomenon corresponds to the behavior of water influx seen during the course of the simulations. Both C36 and Drude gave rise to a similar degree of hydration over each umbrella sampling window, with Drude showing slightly lower hydration in the bilayer, and slightly higher hydration outside of the bilayer (Supporting Information, Figure S7E). The polarization within the bilayer relative to bulk water is due to interactions with phosphate groups that invaginate slightly (Figure 12), helping explain the slightly higher electric field values within the bilayer compared to outside of the bilayer.

**Figure 11.**
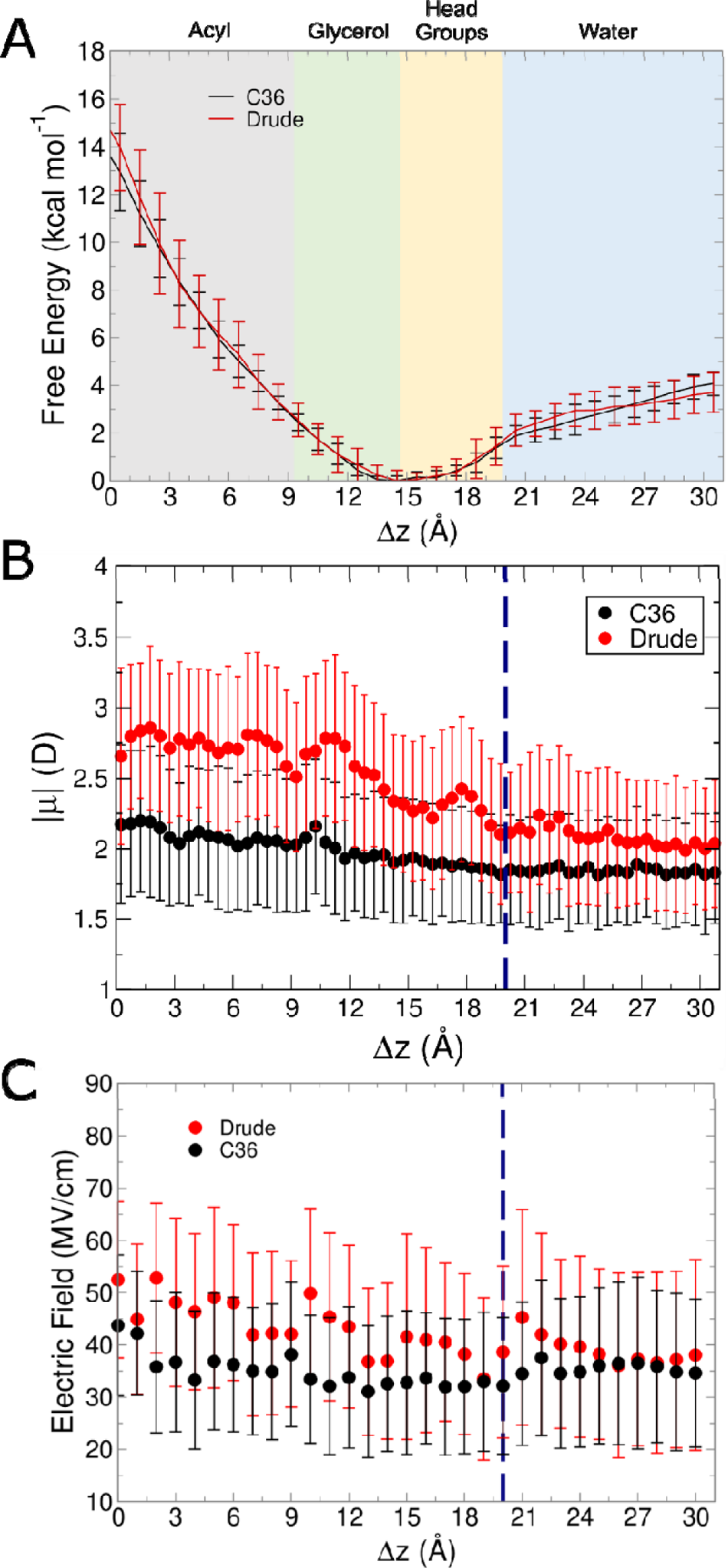
Results from C36 and Drude simulations of methylguanidinium systems. (A) Free energy surfaces from umbrella sampling simulations. (B) Molecular dipole moments as a function of position within the membrane in unbiased simulations. Error bars correspond to the RMSF of the binned data points. (C) Average intrinsic electric field magnitude acting at the COG of methylguanidinium in each umbrella sampling window. Blue dashed lines in panels (B) and (C) denote the average position of the phosphate groups.

**Figure 10.**
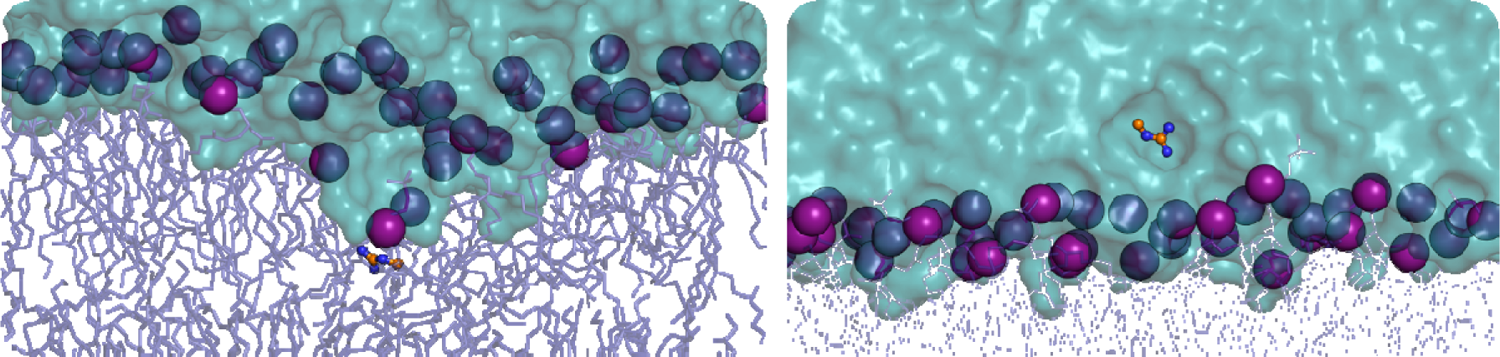
Representative snapshots from Drude simulations of methylguanidinum at Δz = 2 Å (left, at 45 ns) and at Δz = 26 (right, at 45 ns). The methylguanidinum molecule is represented as orange ball-and-stick, lipids tails as dar blue sticks, phosphorus atoms as purple spheres, and water as cyan surface.

Methylguanidinum is one of the better studied amino-acid sidechain analogs, including polarizable FF simulations at an air/water interface^80^ and through umbrella sampling membrane permeation studies.^27^ In both cases, comparisons were made to a nonpolarizable FF. Zhu and Huang found that using a polarizable FF impacted interfacial properties, such as orientation and induced dipole moments, of methylguanidinum at the air/water interface, with corresponding differences in both properties which were in better agreement with experiment using the Drude FF than C36. In addition, Chen *et al.* simulated methylguanidinum with lipids of varying chain length, cholesterol, and mixed lipid systems, performing similar umbrella sampling simulations.^27^ Their results showed a difference in ion transduction mechanism dependent on the FF being used, and dependent on lipid tail length, as we discussed above in the case of acetate. They suggested a transition from ion-induced defect mechanism to solubility diffusion mechanism at a hydrophobic thickness of at least 29 Å for methylguanidinum. POPC has a slightly smaller hydrophobic thickness, resulting in our simulation following the ion-induced defect mechanism for methylguanidinum, as seen by the perturbation of the bilayer in Figure 12. However, for acetate, we observed the ion translocation mechanism depending on the FF used. The positively charged methylguanidinum more readily recruits phosphate groups when permeating across the interface, resulting in the membrane perturbation seen during those simulations. For a bilayer with a thickness comparable to POPC, the cost of deforming the bilayer likely overcomes the cost of dehydrating methylguanidinum. However, for acetate, the cost of permeation, even with a few water molecules, is less than the cost of deforming the bilayer when using the Drude FF. The net result is still largely unfavorable but results in a barrier similar to the cost of methylguanidinum entering the bilayer.

## CONCLUSIONS

Here, we investigated the impact of explicit polarization on the energetics and dynamics of several analogs of amino-acid sidechains. In terms of molecular dipole moment, the Drude FF shows sensitivity to the localization of the molecule in the bilayer, generally with a larger dipole moment toward lipid headgroup regions and a smaller dipole moment within the bilayer interior. In contrast, simulations performed with the C36 FF showed no such sensitivity, as expected. The only variations in dipole moment in these simulations arose from geometric changes and bond vibrations.

When comparing free energy surfaces, in the case of nonpolar, uncharged molecules and positively charged methylguanidinium, results between the two FFs were similar, suggesting that simple additive FFs may represent these species well. Explicit polarization led to differences between the free energy surfaces in aromatic molecules, even in the case of benzene, a relatively simple and symmetric molecule, suggesting that modeling the π system in these species is sensitive to electronic polarization, as expected. In cases of molecules with greater polarity, we found differences in features of the free energy surfaces, with minima shifting, and energy barriers changing. The most striking example of this property is the negatively charged acetate, which differed in the apparent mechanism of ion translocation as a function of FF. Careful analysis of hydration of this molecule showed that the C36 FF likely leads to dramatic over-hydration of the anion when it resides in the membrane core. The Drude FF suggests that some hydration is possible, but to a lesser extent.

Even small differences in free energy barriers, or slight perturbations to preferred localization of a given molecule, will likely have a profound impact when summed over the tens or hundreds of residues in a membrane protein. The energetic consequences of explicitly treating polarization may therefore impact the preferences and dynamics of individual residues, especially in cases of amino acids buried in heterogeneous microenvironments or those at the interface, which may lead to shifts in the dynamics of the protein. In addition, using a polarizable FF provides a finer detail of the underlying electrostatic forces driving dynamics and electrostatic gradient of the bilayer, which may be lacking with nonpolarizable FFs. As such, the use of polarizable FFs such as the classical Drude oscillator model represents a promising step forward in simulating membrane and membrane-protein systems.

## ACKNOWLEDGMENTS

The authors thank Virginia Tech Advanced Research Computing for providing computing resources and support. This work also used the Expanse Cluster at San Diego Supercomputer Center (SDSC) through allocation BIO220106 (to JMM) and allocation BIO230117 (to JAL) from the Advanced Cyberinfrastructure Coordination Ecosystem: Services & Support (ACCESS) program,^81^ which is supported by National Science Foundation grants #2138259, #2138286, #2138307, #2137603, and #2138296. This material is based upon work supported by the National Science Foundation Graduate Research Fellowship under grant #1840995 (to JMM), by the National Institutes of Health (grant R35GM133754 to JAL), and the U.S. Department of Agriculture National Institute of Food and Agriculture (project number VA-160092 to JAL).

## SUPPORTING INFORMATION AVAILABLE

A PDF file containing 1 table and 6 figures including total simulation times per system for umbrella sampling, data for unbiased simulations of indole and methylimidazole from alternate starting orientations, analysis of hydration and hydrogen bonding for the umbrella sampling simulations, and convergence checks of the free energy surface of the umbrella sampling simulations.

## TOC Graphic

For Table of Contents Only

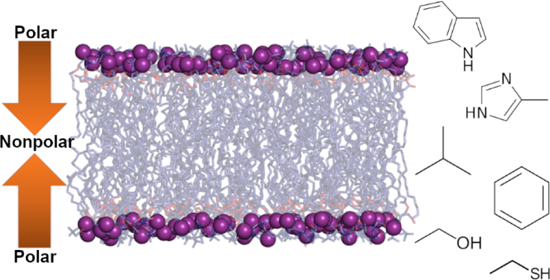

