## Supporting Information for "Quantifying Induced Dipole Effects in Small Molecule Permeation in a Model Phospholipid Bilayer"

**SUPPORTING TABLES**

**Table S1.** Simulation times (ns) for umbrella sampling windows in each system.

|  | **CHARMM36** | **Drude** |
| --- | --- | --- |
| Isobutane | 50 | 70 |
| Benzene | 80 | 50 |
| Indole | 50 | 90 |
| Methylimidazole | 50 | 50 |
| Ethanethiol | 50 | 50 |
| Ethanol | 50 | 80 |
| Acetate | 50 | 70 |
| Methylguanidinium | 50 | 50 |

**SUPPORTING FIGURES**

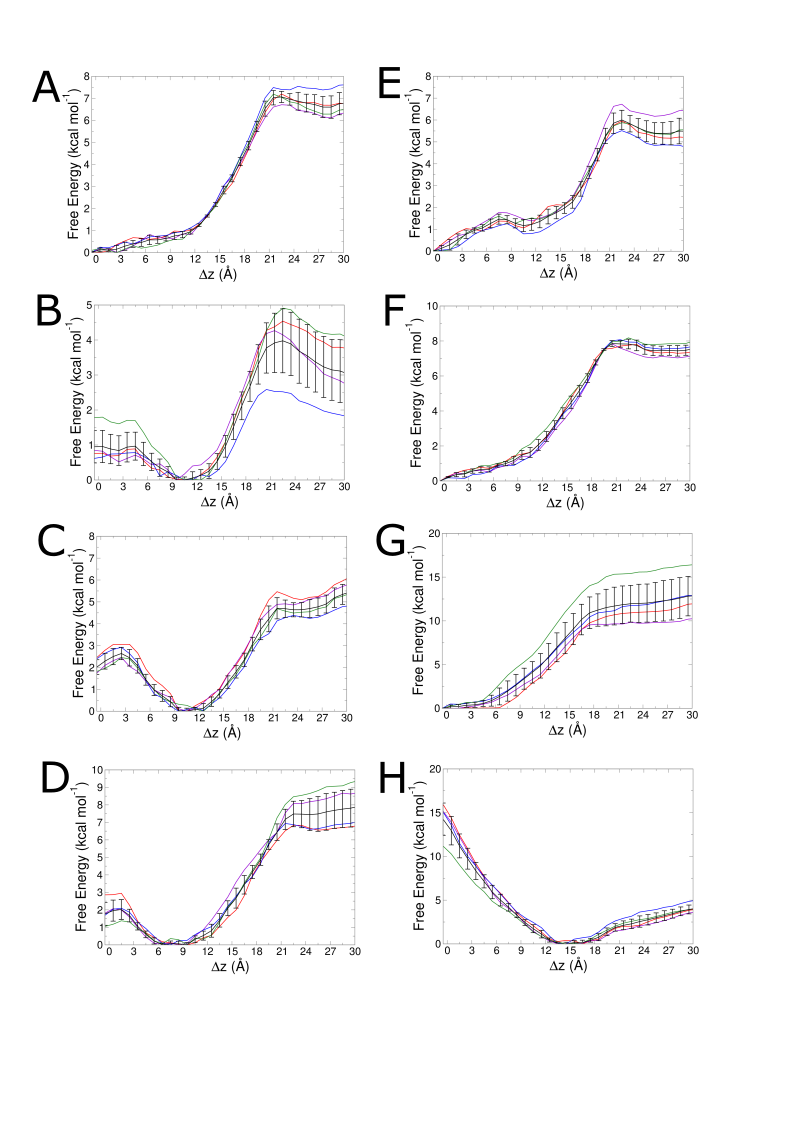

**Figure S1.** Convergence checks for C36 simulations. Free energy surfaces from last 40 ns in 10 ns intervals are plotted in different colors (in order: green, blue, purple, red) along with the pooled data (black with error bars) for (A) isobutane, (B) benzene, (C) indole, (D) methylimidazole, (E) ethanethiol, (F) ethanol, (G) acetate, (F) methylguanidinum.

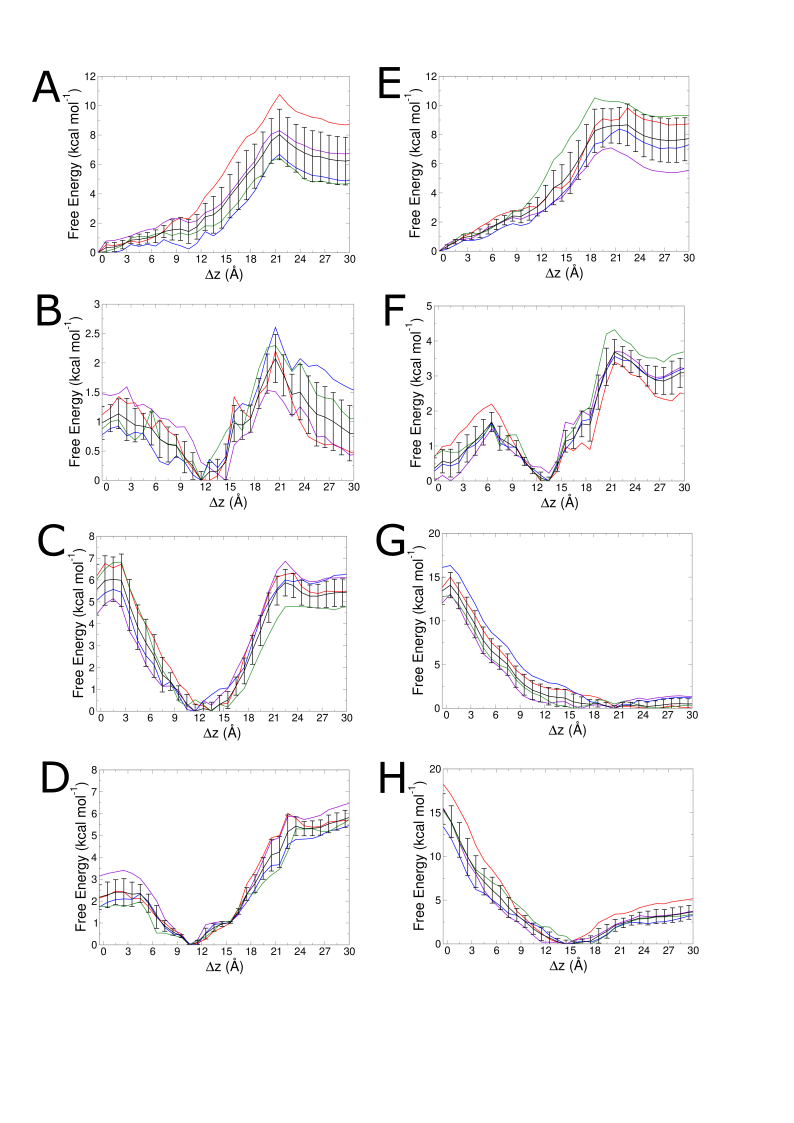

**Figure S2.** Convergence checks for Drude simulations. Free energy surfaces from last 40 ns in 10 ns intervals are plotted in different colors (in order: green, blue, purple, red) along with the pooled data (black with error bars) for (A) isobutane, (B) benzene, (C) indole, (D) methylimidazole, (E) ethanethiol, (F) ethanol, (G) acetate, (F) methylguanidinum.

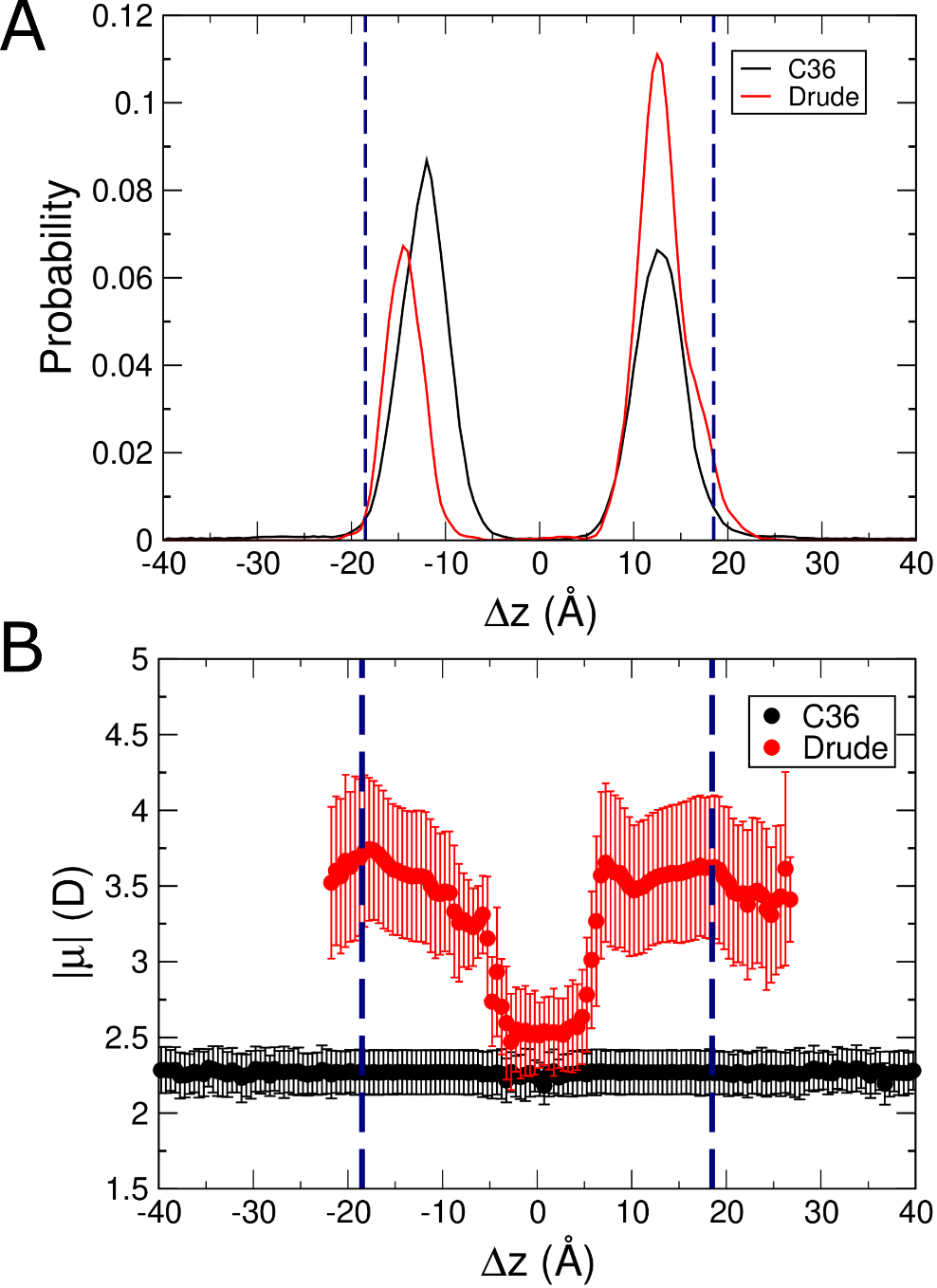

**Figure S3.** Results from unbiased simulations of indole starting from the flipped orientation. (A) Normalized probability of localization of indole in unbiased simulations. (B) Molecular dipole moments as a function of position within the membrane in unbiased simulations. Error bars correspond to the root-mean-squared fluctuation (RMSF) of the binned data points.

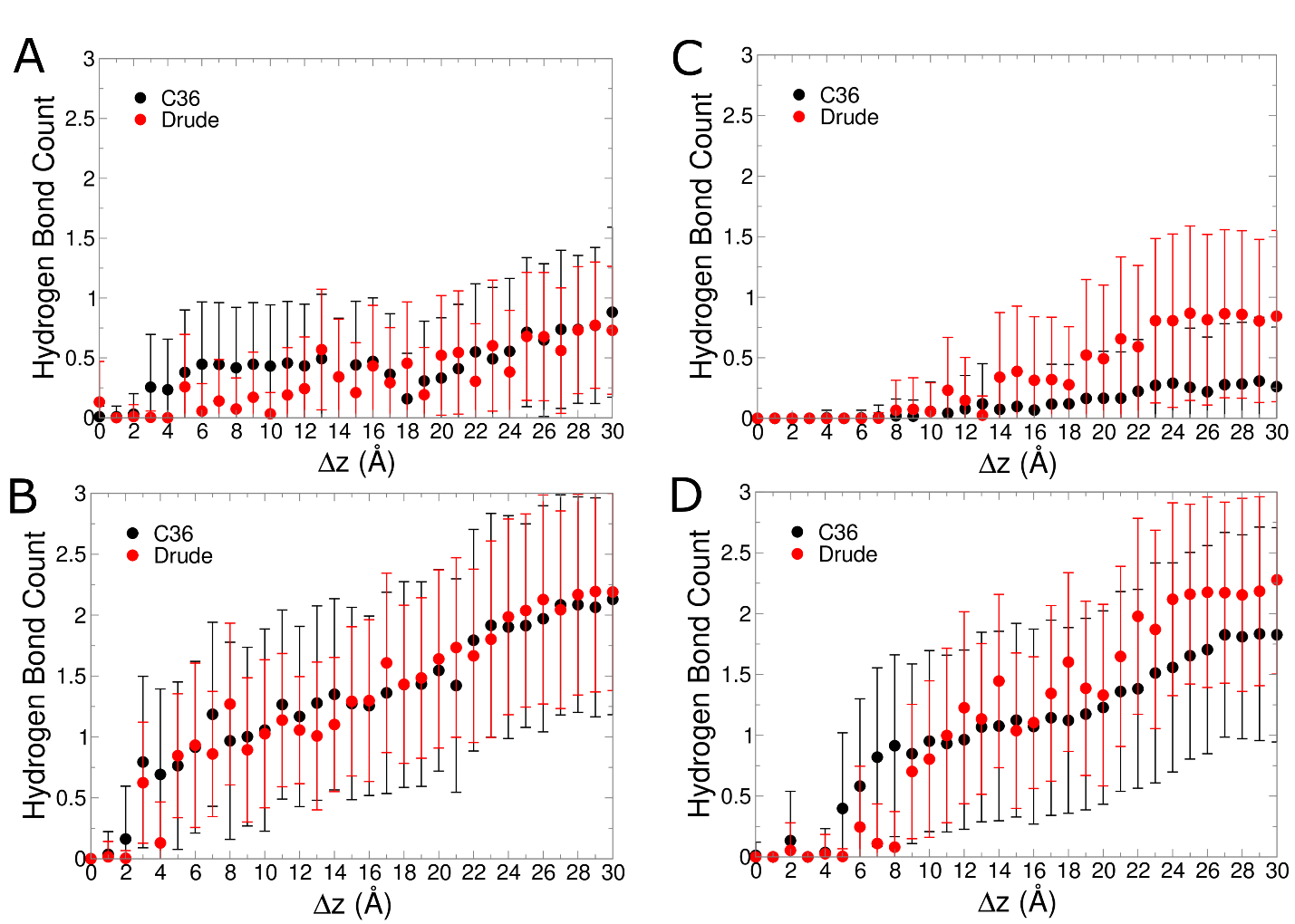

**Figure S4.** The average number of hydrogen bonds occurring between water and the small molecule in each umbrella sampling window. (A) Indole, (B) methylimidazole, (C) ethanethiol, and (D) ethanol. Error bars correspond to the root-mean-squared fluctuation (RMSF) of the binned data points.

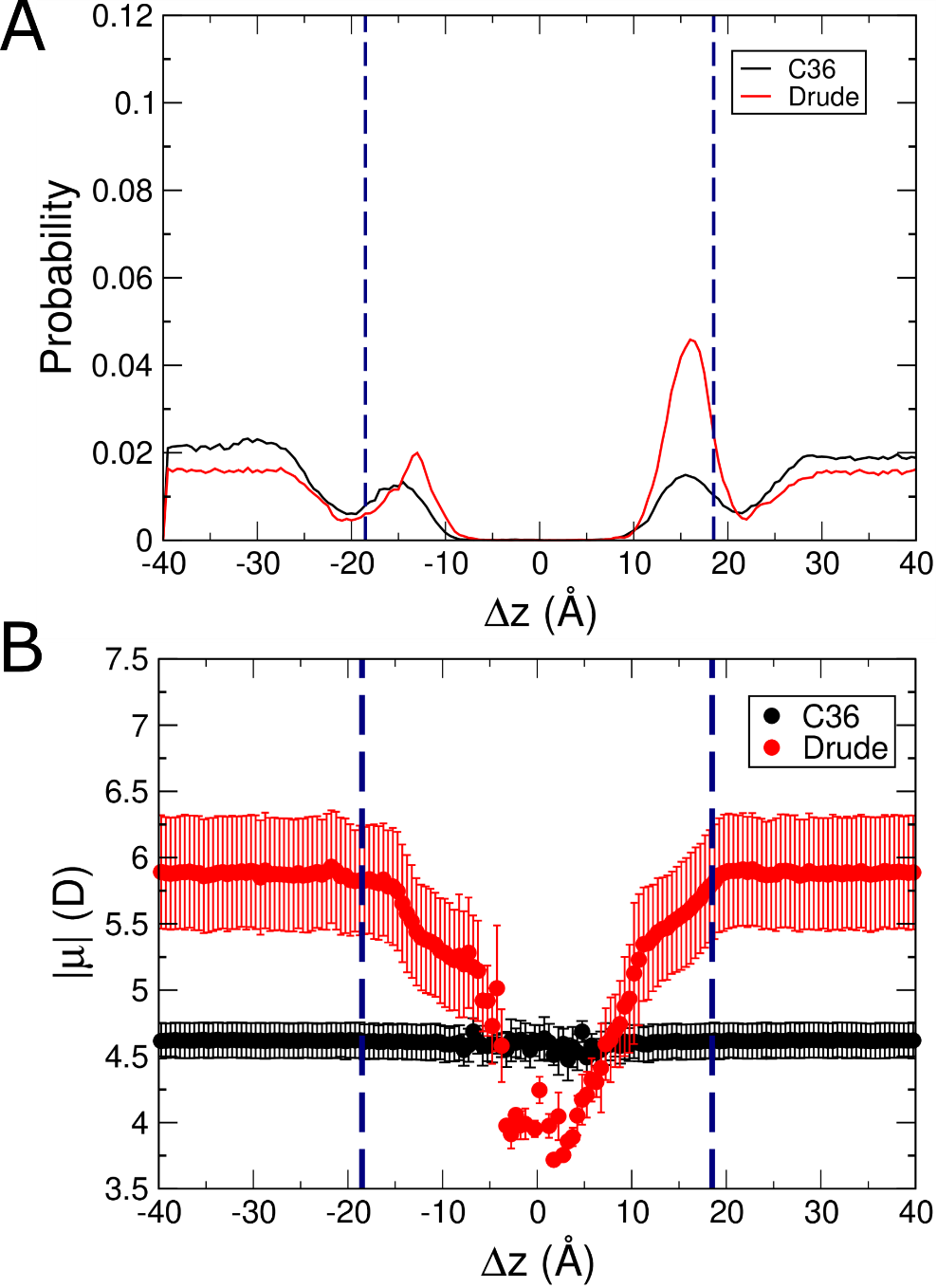

**Figure S5.** Results from alternate unbiased simulations of methylimidazole starting from the flipped orientation. (A) Normalized probability of localization of methylimidazole in unbiased simulations. (B) Molecular dipole moments as a function of position within the membrane in unbiased simulations. Error bars correspond to the RMSF of the binned data points.

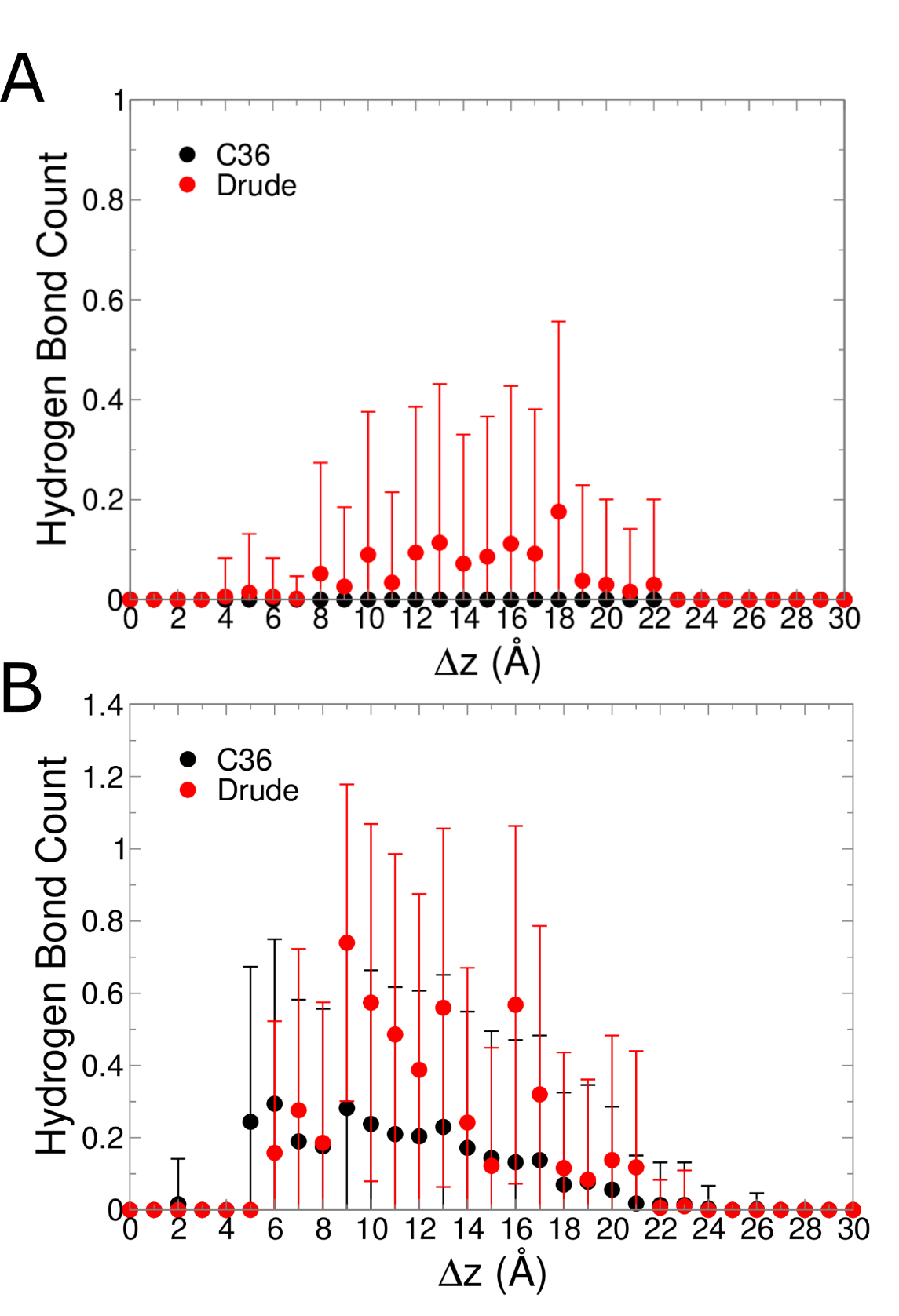

**Figure S6.** The average number of hydrogen bonds formed between POPC and the small molecule in each umbrella sampling window. (A) Ethanethiol, and (B) ethanol. Error bars correspond to the root-mean-squared fluctuation (RMSF) of the binned data points.

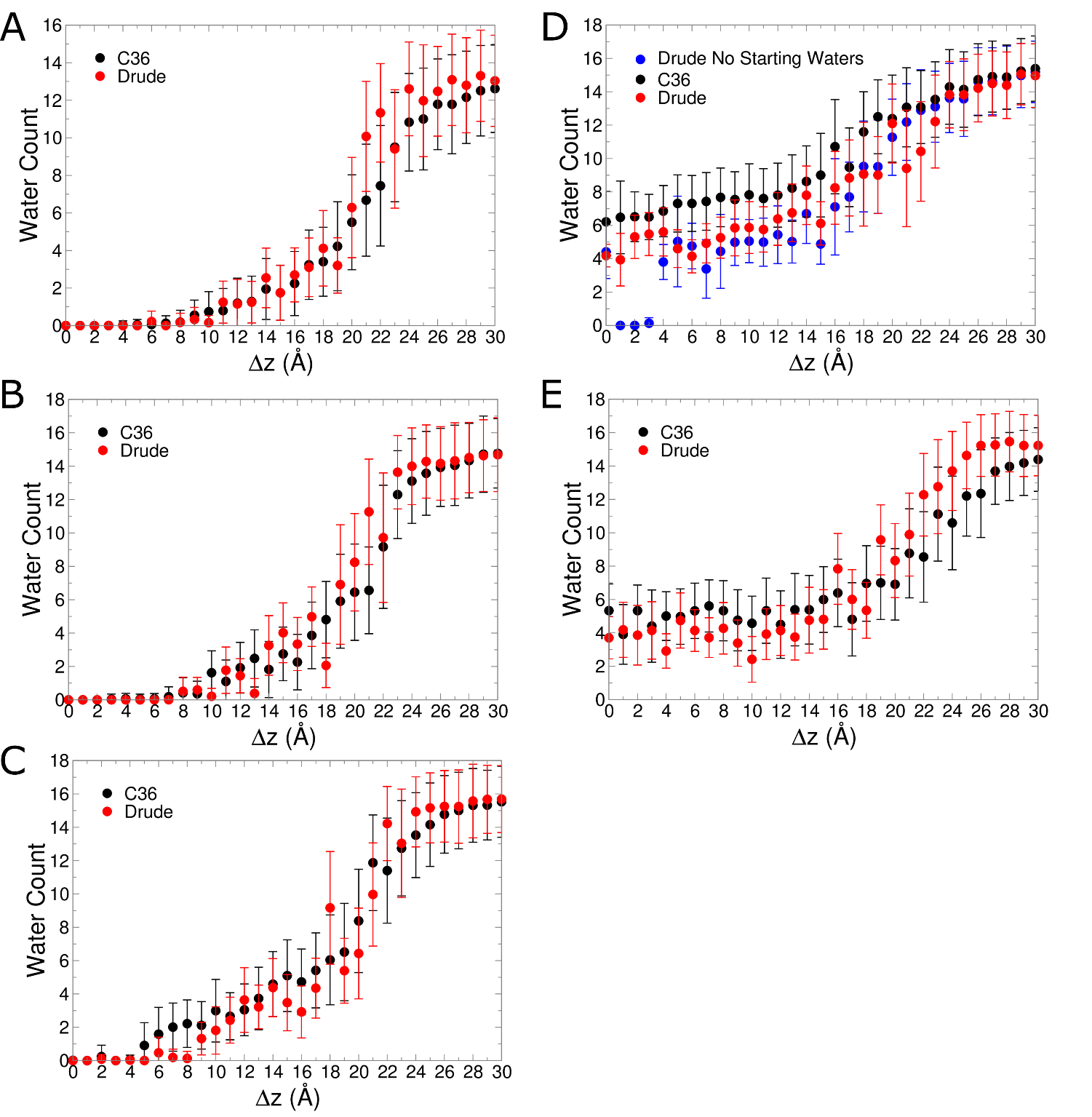

**Figure S7.** The average number of water molecules within 5 Å of the center of mass of each small molecule in each umbrella sampling window. (A) Benzene, (B) ethanethiol, (C) ethanol, (D) acetate (including results from the alternate starting coordinates), and (E) methylguanidinum. Error bars correspond to the root-mean-squared fluctuation (RMSF) of the binned data points.
